# Arthritis Flares Mediated by Tissue Resident Memory T Cells in the Joint

**DOI:** 10.1101/2021.06.04.446927

**Authors:** Margaret H Chang, Anaïs Levescot, Nathan Nelson-Maney, Rachel B Blaustein, Kellen D Winden, Allyn Morris, Alexandra Wactor, Spoorthi Balu, Ricardo Grieshaber-Bouyer, Kevin Wei, Lauren A Henderson, Yoichiro Iwakura, Rachael A Clark, Deepak A Rao, Robert C Fuhlbrigge, Peter A Nigrovic

## Abstract

Although rheumatoid arthritis is a systemic disease, flares typically occur in a subset of joints that is distinctive for each patient. Pursuing this intriguing pattern, we show that arthritis recurrence is mediated by long-lived synovial resident memory T cells (T_RM_). In three murine models, CD8+ cells bearing T_RM_ markers remain in previously inflamed joints during remission. These cells are bona fide T_RM_, exhibiting failure to migrate from joint to joint, preferential uptake of fatty acids, and long-term residency. Disease flares result from T_RM_ activation by antigen, leading to CCL5-mediated recruitment of circulating effector cells. Correspondingly, T_RM_ depletion ameliorates recurrence in a site-specific manner. Human rheumatoid arthritis joint tissues contain a comparable CD8+-predominant T_RM_ population, most evident in late-stage non-inflamed synovium, exhibiting limited T cell receptor diversity and a pro-inflammatory transcriptomic signature. Together, these findings establish synovial T_RM_ cells as a targetable mediator of disease chronicity in autoimmune arthritis.

## Introduction

Rheumatoid arthritis (RA) affects millions of individuals worldwide (1). These chronic autoimmune disorders are characterized by immune-mediated inflammation within joint tissue (2). While arthritis can in many cases improve with immunomodulatory therapies, treatment is often life-long, and episodic flares of joint inflammation continue to affect most patients (3).

Arthritis flares exhibit a strong tendency to recur in a restricted subset of joints. In RA, most joints that will ever be affected become inflamed within the first year of disease; subsequent involvement of a previously spared joint is unusual (4). This finding highlights the presence of a mechanism mediating local memory, but the nature of this mechanism remains unexplored.

Memory T cells are immune cells that develop after antigen exposure and persist for a lifetime. Resident memory T cells (T_RM_) develop in tissues after antigen exposure, taking up long-term residence to provide local site-specific immune defense against pathogens previously encountered in these locations (5–8). Unlike other memory T cells, T_RM_ do not recirculate via the blood or lymphatics (9, 10). Belonging to either CD4 or CD8 T cell subsets, T_RM_ have been described in the lung, gut, brain, reproductive tract and skin, but not in synovial tissue (6–8, 11–13). They exhibit effector functions such as cytokine production and cytolysis, activation of local innate and adaptive immune cells, and recruitment of other effector cells from the circulation (11, 14–16). T_RM_ contribute to local immune defense and tissue homeostasis (9, 11), but they can also drive localized recurrent inflammation; for example, psoriasis and fixed drug eruptions are both mediated by T_RM_ (5, 17–19). We sought to investigate a role for T_RM_ in recurrent sitespecific joint inflammation.

Using three murine models of arthritis, we show long-term persistence of CD8+ T_RM_ in arthritic joints during remission. Activation of these cells by antigen led to recruitment of circulating lymphocytes and arthritis flare. Site-specific depletion during remission curtailed arthritis recurrence, demonstrating an essential role for T_RM_ within quiescent joints in instigating recurrent, joint-specific flares. Correspondingly, we identified T_RM_ cells in human RA synovial tissue, most prominently in late-stage noninflamed synovium. These T_RM_ are primarily CD8+ and exhibit limited T cell receptor diversity, as well as transcriptional profiles consistent with immune activation and cell recruitment that support a pro-inflammatory role. Together, these findings provide a mechanistic explanation for joint-restricted disease recurrence in autoimmune arthritis.

## Results

### Generation of a murine model of recurrent synovitis

To study arthritis flare, we developed an animal model of recurrent synovitis by adapting a T cell-driven, joint-specific model of antigen-induced arthritis (Figure 1A) (20–22). Arthritis was established by intraarticular injection of methylated bovine serum albumin (meBSA) without adjuvant into one wrist, knee and ankle of a B6 mouse, while contralateral joints were injected with vehicle (phosphate-buffered saline, PBS) as an internal control. Mice also received IL-1β injected into the footpad on either side on days 0-2 (20, 22). By 1 week, mice developed joint swelling restricted to the antigen-exposed joints (Figure 1B), quantified by measurement of the anterior-posterior diameter of the wrist as the joint measured most reliably (Figure 1C). Clinical swelling subsided by 4 weeks without further intervention but could be re-activated by intraperitoneal (i.p.) injection of meBSA, resulting in recurrent inflammation limited to previously inflamed joints, which we termed a “flare” in this model (Figure 1C-D). Characterization of the joint by histology revealed synovial thickening and infiltration of inflammatory cells in the antigen-exposed joint at week 1, corresponding to acute joint swelling assessed clinically (Figure 1E). Inflammatory infiltrates resolved without intervention by 4 weeks but recurred in previously inflamed joints during the flare response to a systemic antigen challenge (Figure 1E).

**Figure 1.**
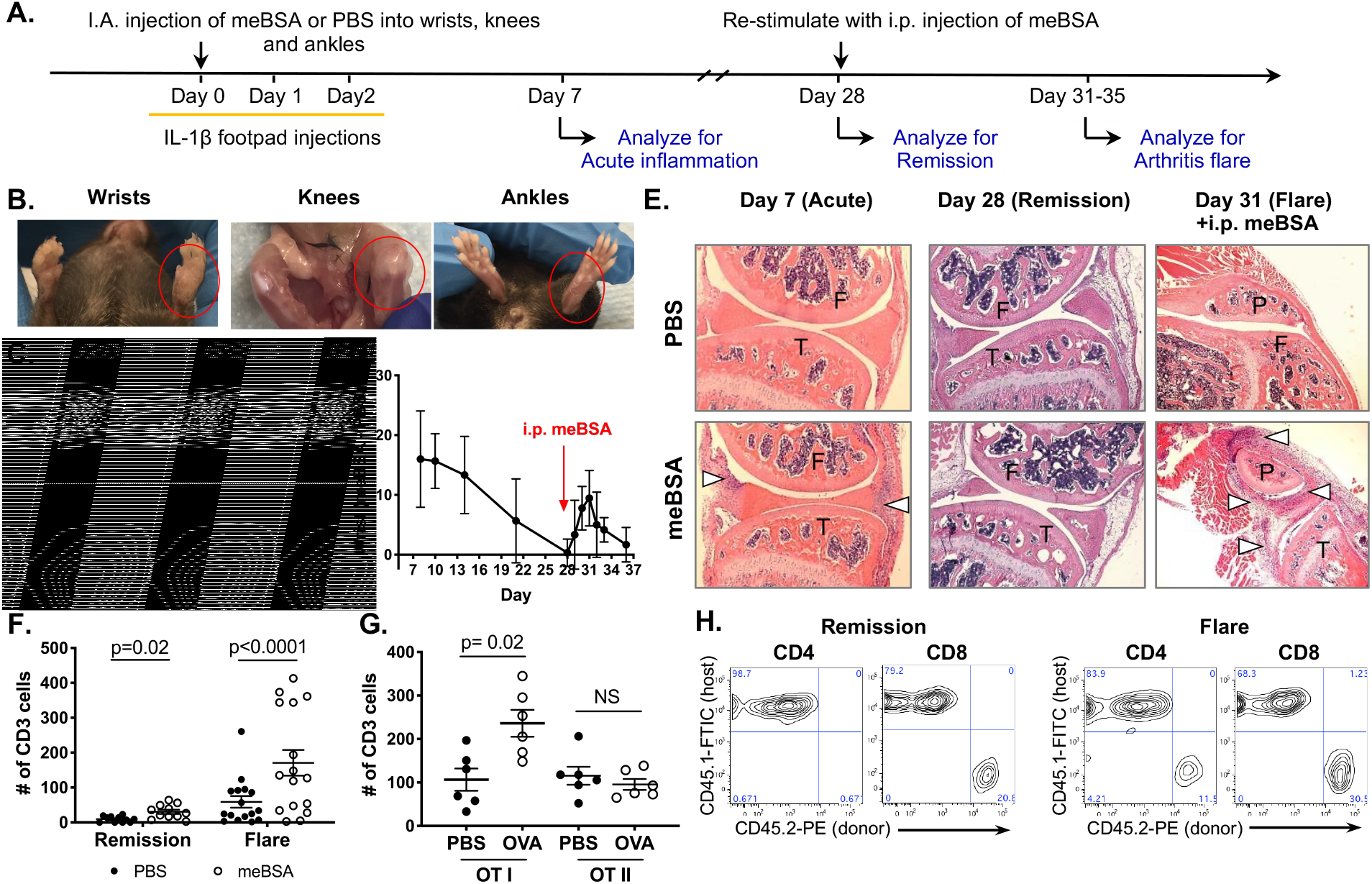
Arthritis flare model identifies CD8 T cells as mediators of recurrent synovitis. **(A)** Diagram of mouse model of arthritis flare. **(B)** Representative images of wrists, knees and ankles of arthritic mouse at Day 7. Red circle denotes antigen-injected joint. **(C)** Measured wrist thickness at Day 7, Day 28 and Day 31. Each dot represents one animal (n=15, day 7 and 28; n=9, day 31). p-values from two-tailed paired student’s t-test. **(D)** Difference in wrist thickness of meBSA-vs. PBS-injected joints over time (n=15 mice). **(E)** Representative H&E of contralateral PBS-vs. meBSA-injected knees from the same mouse at indicated timepoints. (P) patella, (F) femur, (T) tibia, arrows mark inflammatory infiltrates. **(F)** Number of synovial T cells during remission and flare. Each dot represents one animal (n=11, remission; n=16, flare). p-values from two-tailed Wilcoxon matched-pairs signed rank test. **(G)** Synovial T cell count after adoptive transfer of OT-I and OT-II T cells. Each dot represents one animal (n=6 OT-I; n=6 OT-II). p-values from two-tailed paired student’s t-test. **(H)** Representative contour plots of CD45.2+ donor-derived vs. CD45.1+ host-derived CD4 and CD8 T cells in the synovium during remission and flare.

Arthritis mediated by meBSA depends on T cells, as RAG-1^-/-^ mice and CD4-depleted mice fail to develop disease (22). To investigate the immune cells involved in the arthritis flare, we assessed dissociated synovium using flow cytometry three days after systemic antigen re-stimulation (Day 31). We found CD19+ B cells, CD3+ T cells, CD11b+Ly6G-monocyte/macrophage cells, and a large CD11b+Ly6G+ neutrophil population within the synovium of joints previously exposed to antigen; these cells were rare in the contralateral joints not injected with antigen (Supplemental Figure 1). While no clinical swelling or histological evidence of inflammation remained during remission, a population of T cells persisted within the synovium of previously inflamed joints (Figure 1F, remission). After systemic antigen challenge, this population expanded markedly (Figure 1F, flare), supporting a potential role in disease reactivation.

### Antigen-specific synovial CD8 T cells mediate recurrent synovitis

To determine a role for CD4 and CD8 T cells in arthritis flares, we utilized an adoptive transfer approach using donor mice transgenic for ovalbumin (OVA)-specific TCR (OT-I, CD8 T cells; OT-II, CD4 T cells). We collected peripheral lymph nodes from 5-6 week-old CD45.2+ OT-I or OT-II mice, isolated CD3+ cells through negative selection, and transferred CD3+ cells IV into recipient wildtype B6 CD45.1 mice on day - 1. Joints were then injected with OVA (instead of meBSA) on day 0 per Figure 1A, and during remission the mice were challenged with i.p. OVA to trigger an arthritis flare. Adoptive transfer of antigen-specific OT-I T cells yielded a flare response, as characterized by T cell infiltrates, while transfer of antigen-specific OT-II T cells did not, indicating dependence of the flare on CD8 T cells (Figure 1G). When OT-I and OT-II T cells were transferred together, all donor (CD45.2+) T cells remaining in the synovium during remission were CD8+, while both CD4+ and CD8+ donor T cells were recruited back to the synovium during a flare (Figure 1H).

To determine whether the arthritis flare is antigen-specific, we generated arthritis with intra-articular meBSA and challenged with i.p. injection of a naïve antigen (OVA) or non-specific inflammatory trigger such as inflammatory cytokines (IL-1β) or TLR ligands (LPS) (Supplemental Figure 2A). Only re-stimulation with the original antigen (meBSA) was able to produce an arthritis flare in the previously inflamed joint (Supplemental Figure 2B). Extending this line of investigation, we generated arthritis with intra-articular meBSA and introduced OT-I T cells during remission to confer OVA reactivity. We then challenged the mice with i.p. injection of meBSA or OVA (Supplemental Figure 2C). The availability of circulating OVA-specific T cells was insufficient to trigger joint disease as joint inflammation occurred only with meBSA re-stimulation, further confirming the key role of antigen-specific T cells remaining within the joint after the initial arthritis episode (Supplemental Figure 2D-E).

These data indicate that antigen-specific CD8 T cells, recruited into the synovium during the initial inflammatory response and retained during remission, participate in joint-specific arthritis recurrence.

### A distinct population of memory T cells with T_RM_ phenotype is present in previously inflamed joints

To identify the T cells involved in joint-specific memory, we assessed memory cell markers on synovial T cells from previously inflamed joints. Arthritis was induced by intraarticular OVA or PBS injection into mice that received adoptive transfer of OT-I T cells. After 1 month of remission, flare was triggered by i.p. OVA and synovium was collected 3 days later for flow cytometry analysis. tSNE analysis of CD45+CD3+ T cells showed a distinct population of T cells within OVA-treated joints that was absent from PBS-injected joints (Figure 2A). These cells expressed CD8+CD44+CD62L-CD69+CD103+/-ST2+ surface proteins, consistent with the cellular profile of T_RM_. Strikingly, cells expressing the T_RM_ signature (see Supplemental Figure 3 for gating strategy) remained in the synovium of previously inflamed joints during remission (Figure 2B, left & Figure 2C, remission), despite absence of joint swelling or histological evidence of inflammation (Figure 1C-E). These T_RM_-like cells expanded during an arthritis flare, again consistent with an active role in joint inflammation (Figure 2B, right & Figure 2C, flare).

**Figure 2.**
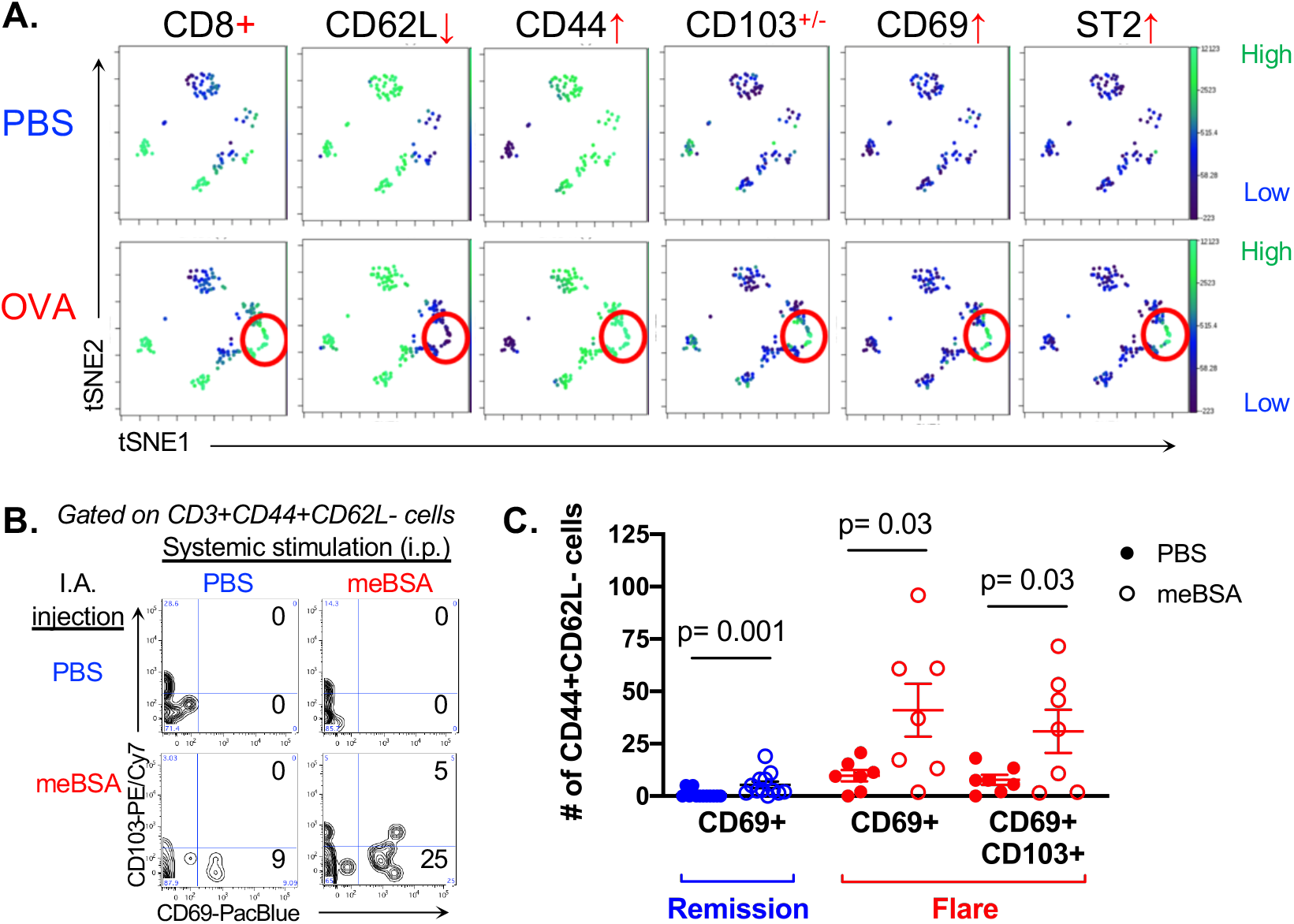
Distinct population of T_RM_-like memory T cells identified in previously inflamed synovium. **(A)** tSNE analysis comparing cellular protein expression in synovial T cells from PBS-injected (top row) vs. OVA-injected (bottom row) joints pooled from 5 mice. Red circle highlights distinct cell population found in OVA-injected joints. **(B)** Representative contour plots comparing CD69 and CD103 expression in CD3+CD44+CD62L-T cells in the synovium of paired PBS-injected (top row) vs. meBSA-injected (bottom row) joints from the same mouse. The left column reflects a mouse in remission while the mouse in the right column was challenged with systemic antigen to induce a flare. **(C)** Quantification of CD69+ and CD69+CD103+ CD44+CD62L-T cells during remission and flare. Each dot represents one animal (n=12, remission; n=7, flare). p-values from two-tailed Wilcoxon matched-pairs signed rank test.

### Synovial memory T cells persist in the synovium and do not migrate in response to tissue egress signals

To confirm that these synovial memory T cells are in fact T_RM_, we assessed their persistence in tissues using two-color fluorescent Cre-reporter mice (mT/mG mice) that express tdTomato at baseline but switch to EGFP after Cre-mediated recombination (23). We induced arthritis in both knees, then injected adeno-associated virus expressing Cre recombinase (AAV-Cre) into a single knee to label the cells remaining in that joint during remission (Figure 3A). Adeno-associated viruses are not replication competent, restricting labelling of cells to a single timepoint (24, 25). One month after AAV-Cre injection, we assessed the synovium for EGFP-positive T cells. We found a distinct population of EGFP+ T cells in joints injected with AAV-Cre that expressed CD8+CD44+CD62L-CD69+, consistent with CD8+ T_RM_ (Figure 3B-C). Importantly, no EGFP+ T cells were found in the contralateral knee, confirming that synovial T cells remained anchored within the previously inflamed joint.

**Figure 3.**
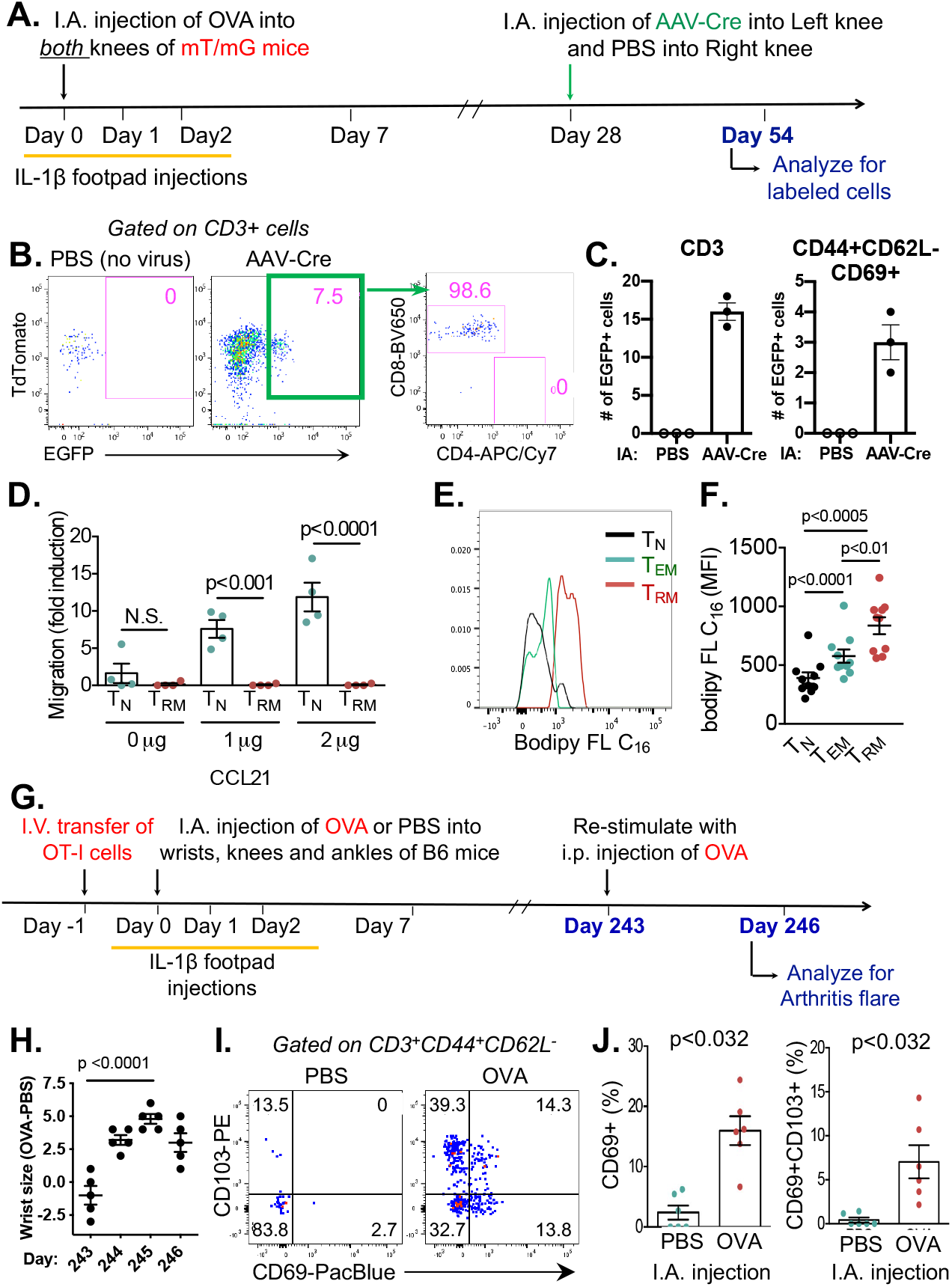
Synovial memory T cells are true T_RM_. **(A)** Experimental design for tracking cells from a specific joint. **(B)** Representative dot plots of T cells pooled from the synovium of 4 mice at Day 54. AAV-Cre-injected and contralateral PBS-injected knees are depicted in paired dot plots with subanalysis of CD8 and CD4 expression in AAV-Cre-labeled T cells (EGFP+). **(C)** Number of AAV-Cre-labelled EGFP+ T cells (*left*) and percentage of CD44+CD62L-CD69+ T_RM_ within EGFP+ T cells (*right*) at Day 54. Each dot represents an independent experiment pooling synovium from 4 mice (n=3). **(D)** Proportion of synovial naïve T cells (T_N_, CD44-CD62L+) and resident memory T cells (T_RM_, CD44+CD62L-CD69+) migrating to increasing concentration of CCL21 chemoattractant. Values displayed as fold change of cells migrated to the bottom compared to cells on top of the transwell. Each dot represents an independent experiment pooling synovium from 5 animals (n=4). p-value from one-way ANOVA. **(E)** Representative histogram of Bodipy FL C_16_ fatty acid uptake by naïve T cells (T_N_), effector memory T cells (T_EM_, CD3+C44+CD62L-CD69-) and resident memory T cells (T_RM_) in synovium. **(F)** Bodipy FL C_16_ fatty acid uptake, depicted as mean fluorescence intensity. Each dot represents one animal (n=10). p-value calculated from one-way ANOVA. **(G)** Experimental design for assessing T_RM_ longevity. **(H)** Difference in wrist size between OVA- and PBS-injected wrists within the same animal. Each dot represents one animal (n=5). p-value from two-tailed student’s t-test. **(I)** Representative dot plot of CD69 and CD103 expression in CD44+CD62L-T cells from synovium on Day 246. **(J)** Percentage of CD44+CD62L-T cells that are CD69+ or CD69+CD103+ joints at Day 246. Each dot represents one animal (n=6). p-values from two-tailed Wilcoxon matched-pairs signed rank test.

We also tested the capacity of synovial T cells to migrate in response to the tissue egress signal CCL21, a chemokine that binds to CCR7 and attracts T cells to secondary lymphoid organs. Sorted CD3+ T cells isolated from the synovium of mice in an arthritis flare were placed in the top of a transwell chamber and CCL21 chemoattractant was placed in the bottom. After 2 hours, cells in the top and bottom chambers were analyzed by flow cytometry. Consistent with *in vivo* results from the AAV-Cre experiment, synovial T_RM_-like cells (CD3+CD44+CD62L-CD69+) failed to migrate across the transwell membrane, whereas synovial naïve T cells (CD3+CD44-CD62L+) showed the expected dose-dependent migration in response to CCL21 (Figure 3D). Thus T_RM_-like cells from inflamed joints remain sessile both *in vivo* and *in vitro*.

### Synovial T_RM_-like cells display increased fatty acid uptake

Skin T_RM_ preferentially utilize exogenous free fatty acids for metabolism, a process essential to their maintenance, longevity and function (26). To determine whether synovial T_RM_-like cells similarly exhibit enhanced uptake of free fatty acids, we collected synovium from mice in an arthritis flare and incubated disaggregated synovial cells with Bodipy FL C16, a green fluorescent free fatty acid. After 30 minutes, we assessed free fatty acid uptake by each T cell subtype using flow cytometry. As with T_RM_ from the skin, synovial T_RM_-like cells (CD3+CD44+CD62L-CD69+) incorporated higher amounts of free fatty acids compared to naïve T cells (CD3+CD44-CD62L+) and effector memory T cells (CD3+CD44+CD62L-CD69-) (Figure 3E-F).

### Synovial T_RM_ retain long-term capacity to initiate arthritis flares

T_RM_ provide long-term immunity in tissues, but aberrant activity can also lead to enduring autoimmunity (5, 19). To determine the longevity of synovial T_RM_, we utilized the OT-I adoptive transfer method to induce arthritis in mice and waited for over 8 months of remission before challenging mice with systemic antigen (Figure 3G). After 243 days, mice still exhibited a flare response restricted to joints that had previously been exposed to antigen (Figure 3H). Evaluation of synovial T cells showed a continued presence of CD3+CD44+CD62L-CD69+ cells in the antigen-injected joint (Figure 3I-J), demonstrating long-term retention and function of synovial T_RM_. Together, surface signature, metabolic features, tissue retention and long-term persistence allow us to conclude that T_RM_-like T cells from inflamed joints in this murine model are bona fide T_RM_.

### T_RM_ persist in remission in a spontaneous murine model of RA

Whereas meBSA and OVA + OT-I T cell models of arthritis are both mediated by exogenous antigen, in the B6 background, we sought to identify T_RM_ in a spontaneous model within a different genetic background. Balb/c mice lacking the endogenous IL-1 receptor antagonist (IL-1Ra) spontaneously develop an intense T cell-dependent inflammatory arthritis with age (27). We noted that some *IL1rn-/-* mice begin with unilateral joint inflammation before accruing additional inflamed joints over time (Figure 4A-B). Remission at this stage could be induced using recombinant IL-1Ra (anakinra); subsequent withdrawal of treatment resulted in an arthritis flare that preferentially occurred in the previously inflamed joint, a pattern of recurrent joint-specific flare reminiscent of human RA (Figure 4C-F). Evaluation of synovial T cells in these mice showed a population of cells with a T_RM_ signature that persisted during remission (Figure 4G-H). These cells were unaffected by antibody-mediated depletion of circulating T lymphocytes with anti-Thy1.2 antibody, establishing these T_RM_ to be tissue resident (Figure 4I-L). The identification of these cells further supports the generalizability of the T_RM_ mechanism in arthritis.

**Figure 4.**
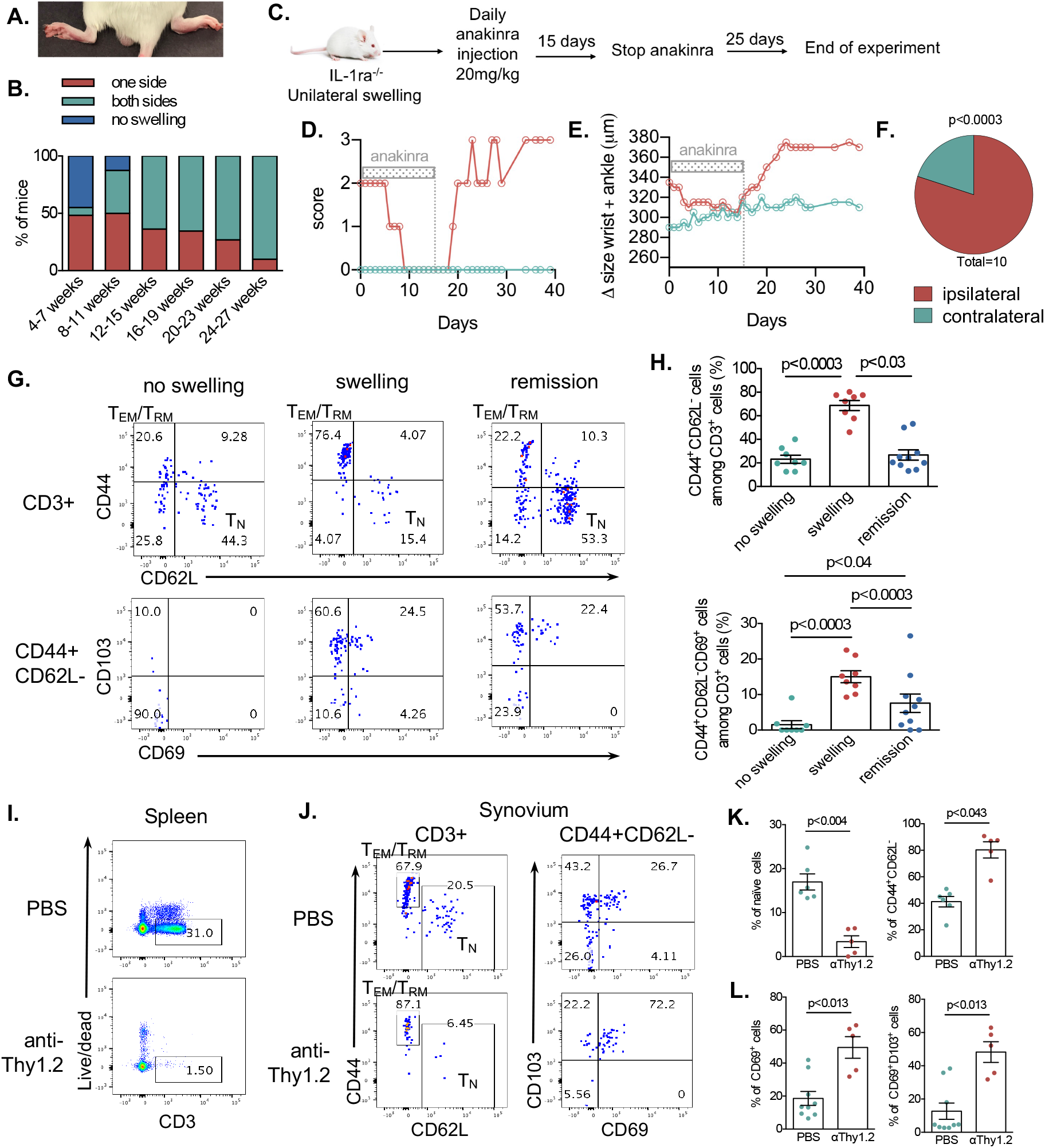
T_RM_ persist in drug-induced remission in spontaneous murine model of RA. **(A)** Representative photo of unilateral arthritis in IL-1 receptor antagonist deficient (*IL-1rn-/-*) Balb/c mice. **(B)** Prevalence of unilateral arthritis by age. **(C)** Experimental design for treating *IL-1rn-/-* mice with unilateral joint swelling with anakinra and subsequent flare after withdrawal of treatment. **(D)** Arthritis score and **(E)** difference in joint measurements of the inflamed joint (red) and contralateral uninflamed joint (green) (n=10). **(F)** Laterality of flare in relation to initial arthritic joint after withdrawal of anakinra. **(G)** Representative dot plots of CD3+ and CD44+CD62L-T cells in the synovium of joints without clinical swelling, with inflammation, and during anakinra-induced remission. **(H)** Graphs quantifying CD44+CD62L-memory T cells and T_RM_ (CD44+CD62L-CD69+) within synovium of the described disease states. (***I-J***) Circulating T cells were depleted in *IL-1rn-/-* mice with i.p. injection of anti-Thy1.2 antibody 250ug 7 days prior to collection. Representative dot plots showing **(I)** anti-Thy1.2 antibody-mediated depletion of CD3+ T cells in the spleen and **(J)** T cell subsets in synovium. Graphs quantifying **(K)** naïve and CD44+CD62L-memory T cells (n=6 PBS, n=5 anti-Thy1.2) and **(L)** CD69+ and CD69+CD103+ memory T cells (n=9 PBS, n=5 anti-Thy1.2) within synovium. Each dot represents one animal. p-values from two-tailed student’s t-test.

### Arthritis flare is dependent on secondary recruitment of circulating lymphocytes

To delineate whether local proliferation of T_RM_ or recruitment of T cells is responsible for the expansion of synovial T cells during flare, we pre-treated arthritic mice in remission with FTY720, a S1P inhibitor which prevents lymphocyte circulation, prior to re-stimulation for flare (Figure 5A). S1P inhibition blocked recurrent joint inflammation, indicating that flare is dependent on secondary recruitment of lymphocytes (Figure 5B-C). Staining for Ki67 expression, a marker of cell proliferation, showed that while there is some local synovial T_RM_ expansion upon antigen re-stimulation (Figure 5D), these cells comprised a negligible fraction of the overall synovial T cell population during arthritis flare (Figure 5E).

**Figure 5.**
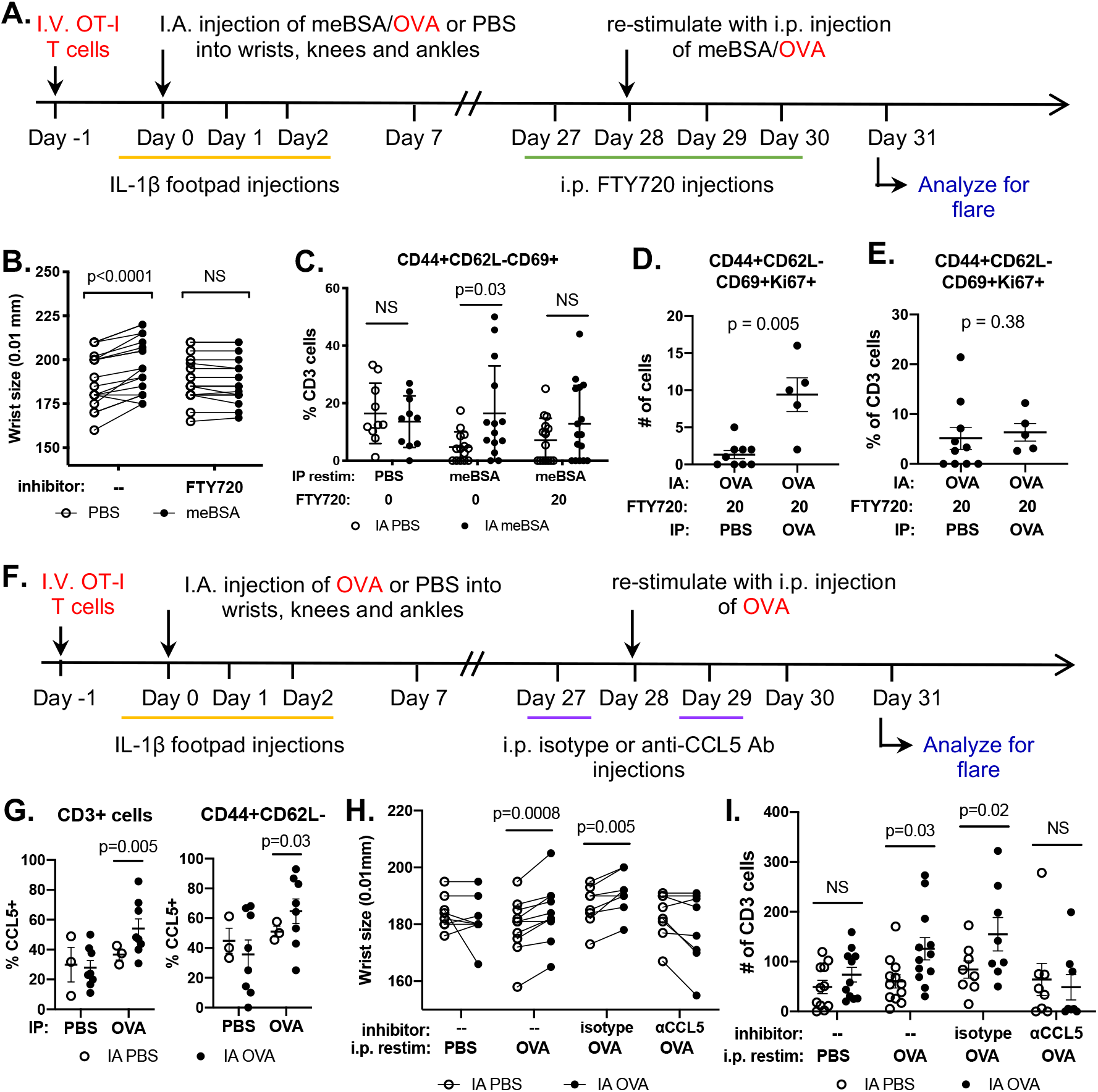
Arthritis flare is dependent on secondary recruitment of circulating lymphocytes. **(A)** Experimental design for inhibiting circulating lymphocytes with FTY720 20ug in meBSA-induced arthritis (black) or with OT-I adoptive transfer and OVA-induced arthritis (red). Graphs depicting **(B)** wrist thickness and **(C)** prevalence of T_RM_ in PBS-vs. meBSA-injected joints at Day 31 flare with FTY720 inhibition. Each dot represents one mouse; line connects contralateral joints within the same animal (n=10 PBS, n=14 meBSA without FTY720, n=16 meBSA with FTY720). p-values from two-tailed paired student’s t-test. **(D)** Number of proliferating Ki67+ T_RM_. **(E)** Ki67+ T_RM_ as a percentage of total CD3+ T cells. Each dot represents one mouse (n=10 pBs, n=5 OVA). p-values from two-tailed Mann-Whitney test. **(F)** Experimental design for inhibiting lymphocyte recruitment with isotype or anti-CCL5 antibody 1ug per dose. **(G)** CCL5 expression in synovial CD3+ cells and CD44+CD62L-T cells during flare assessed by intracellular cytokine staining. Each dot represents one mouse (n=3 IA PBS, n=8 IA OVA). p-values from two-tailed student’s t-test. Graphs depicting **(H)** wrist thickness and **(I)** CD3 T cells in PBS-vs. OVA-injected joints at Day 31 flare with isotype or anti-CCL5 neutralizing antibody. Each dot represents one mouse; line connects contralateral joints within the same animal (n=11 PBS, n=13 OVA without inhibitor, n=8 OVA with isotype or anti-CCL5 antibody). p-values from two-tailed paired student’s t-test.

Chemokines are secreted proteins that act to induce cellular migration during inflammation. CCL5 is prominent in human arthritic synovial fluid, and CD8+CD45RA-T cells have been implicated as major producers of the chemokine (28). We assessed synovial T cells for CCL5, in the absence of exogenous stimulation, and found that some synovial memory T cells contained CCL5 during remission and that this proportion increased markedly during flare (Figure 5G). To investigate the causal contribution of CCL5, we treated mice with an anti-CCL5 neutralizing antibody prior to re-stimulation on days 27 and 29 (Figure 5F). CCL5 inhibition blocked arthritis flare and T cell influx into the joint (Figure 5H-I). These data suggest that T_RM_ induce arthritis flare by recruiting circulating lymphocytes through production of CCL5.

### Synovial resident T cells are required for arthritis flare

To test whether synovial T_RM_ are necessary for arthritis flare, we utilized mice with conditional expression of diphtheria toxin receptor (DTR). Mice expressing Cre-recombinase driven by the *Lck* (lymphocyte protein tyrosine kinase) promoter were crossed with inducible DTR (iDTR) mice that express DTR in the presence of Cre-recombinase. *Lck* is expressed primarily by T cells, associates with CD4 and CD8, and is involved in TCR signaling (29). The resulting Lck-iDTR mice express DTR on all T cells, rendering them susceptible to diphtheria toxin (DT)-mediated depletion. Arthritis was induced in both knees of Lck-iDTR mice (Figure 6A). During remission, DT was injected into one joint while the contralateral knee received intraarticular saline. Intraarticular DT partially depleted synovial T cells (Figure 6B-C). Importantly, intraarticular DT did not deplete circulating T cells, as the percentage of T cells in the peripheral blood remained unchanged during remission, after DT injection, and at restimulation (Supplemental Figure 4). Arthritis flare was then triggered with systemic antigen challenge two weeks after local T cell depletion and joint inflammation was assessed after 72 hours. Localized T cell depletion during remission attenuated arthritis flare as measured by joint histology (Figure 6D-E), T_RM_ expansion, and myeloid cell recruitment (Figure 6F-I, selected as a marker of inflammation to avoid confounding findings resulting from DT-mediated lymphocyte depletion), demonstrating an essential role of resident synovial T cells within quiescent joints in instigating recurrent jointspecific flares.

**Figure 6.**
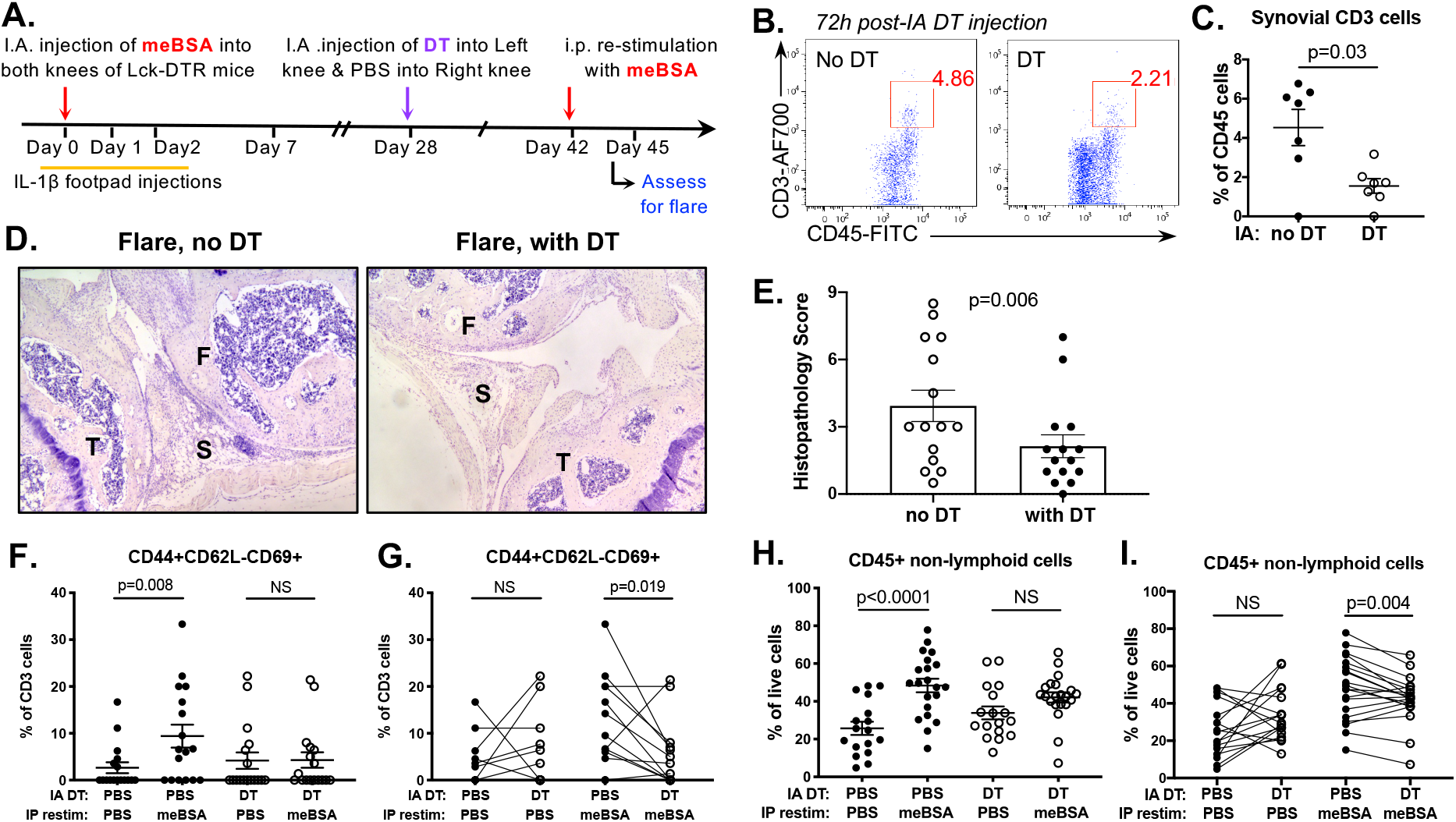
Depletion of synovial resident T cells in remission abrogates arthritis flare. **(A)** Experimental design for localized synovial T cell depletion. **(B)** Representative dot plot of synovial lymphocytes from DT-injected and PBS-injected knees collected 72 hours after intraarticular injection. **(C)** Graph quantifying CD3+ T cells as a percentage of total CD45+ lymphocytes in synovium of DT-treated vs. non-treated knees. p-value calculated from two-tailed paired student’s t-test. Each dot represents one animal (n=6 mice). **(D)** Representative H&E images and **(E)** inflammatory score of contralateral knees from the same mouse with or without DT injection after i.p. re-stimulation for flare at Day 45. Each dot represents one animal (n=15). p-value from two-tailed Wilcoxon matched-pairs signed rank test. (F) femur, (T) tibia, (S) synovium. (***F-I***) Graphs showing percentage of T_RM_ or non-lymphoid CD45+ cells in the synovium in meBSA-injected joints with or without intraarticular DT treatment. **(F,H)** Data compares the flare response with and without DT treatment. **(G,I)** Data evaluates the effect of DT-injection compared to the contralateral control joint. p-values calculated from two-tailed paired student’s t-test. Each dot represents one animal (n=17 mice per condition). Line connects contralateral joints within the same mouse.

### A subset of T cells within human RA synovial tissue displays T_RM_ markers

To determine whether T_RM_ contribute to human RA, we sought T cells with a resident memory signature in human RA synovium. Mantra multispectral immunofluorescence imaging, a tyramide-based immunostaining technology, was performed on formalin-fixed paraffin-embedded synovium obtained from four RA patients. Tissues were stained with antibodies against CD3, CD8, CD4, CD45RO, CD69 and CD103. CD8+ or CD4+ memory (CD3+CD45RO+) T cells were evaluated for coexpression of CD69 and CD103, the most commonly used T_RM_ markers (6–8). We found cells bearing a CD3+CD45RO+CD69+CD103+ signature, consistent with T_RM_, within areas of aggregated lymphoid cells in RA synovium (Figure 7A).

**Figure 7.**
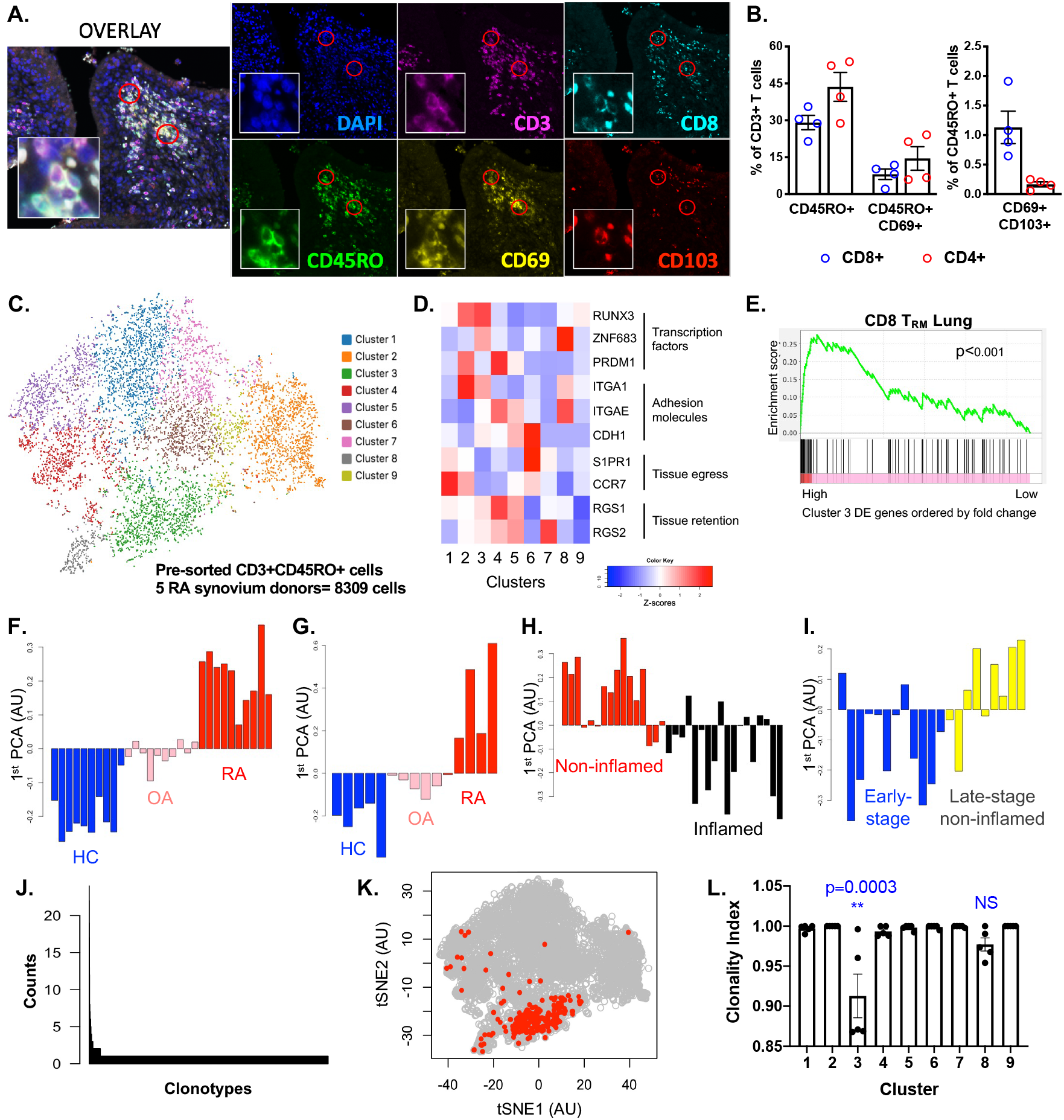
Oligoclonal CD8 T_RM_ enriched in late-stage, non-inflamed RA synovial tissue. **(A)** Representative immunofluorescence image of human RA synovium co-stained for CD3, CD8, CD45RO, CD69 and CD103 proteins and DAPI nuclear stain. Red circle indicates cells positive for all 5 biomarkers. Insert magnifies a circled cell. **(B)** Prevalence of CD8 and CD4 T cells expressing CD45RO memory, CD69 and CD103 markers. Each dot represents the average cell count from 5 fields from each donor (n=4). **(C)** tSNE plot of CD3+CD45RO+ cells from disaggregated RA synovium pooled from 5 donors. Each dot represents 1 cell. Color-coded clusters designate cells with similar gene expression profiles. **(D)** Gene expression heatmap of T_RM_ associated genes. Red=increased expression; blue=decreased expression. **(E)** Gene set enrichment analysis of Cluster 3 differentially expressed (DE) genes to CD8 lung T_RM_ gene expression signature (Kumar et al., 2017). ***(F-G)*** Principal component analysis (PCA) of Cluster 3 DE genes in microarray expression data from healthy controls (HC), osteoarthritis (OA) and rheumatoid arthritis (RA) synovium published in **(F)** Woetzel et al., 2014 (n=30) and **(G)** Ungethuem et al., 2010 (n=15). Each bar represents one donor. ***(H-I)*** PCA of Cluster 3 DE genes in T lymphocytes from RA synovial tissue (Zhang et al, 2019). Bar plots showing Cluster 3 T_RM_ gene signature in **(H)** non-inflamed vs. inflamed RA synovium and **(I)** early-stage RA vs. late-stage non-inflamed RA synovium. Each bar represents one donor (n=34). **(J)** Histogram of cells with the same TCR-alpha and -beta CDR3 sequence. **(K)** Top 0.5% of most frequent T cell clones overlaid on the tSNE plot. **(L)** Clonality index for each cell cluster. Each dot represents one donor (n=5). p-values from one-way ANOVA.

To quantify these cells, we performed unbiased automated identification and analysis of staining within synovial sections. 29.1% (± SEM 2.9) of T cells within human RA synovium were CD8+ memory T cells while 43.6% (± SEM 5.9) were CD4+ memory T cells (Figure 7B). 8.1% (± SEM 2.1) and 14.5% (± SEM 4.8) were CD69+ CD8+ and CD4+ memory T cells. Despite the numeric predominance of CD4+ cells overall, CD69+CD103+ cells were largely CD8+ rather than CD4+ (1.13% ± SEM 0.27 vs. 0.16% ± SEM 0.04 of all memory T cells, respectively) (Figure 7B). Lymphocytes with the CD3+CD45RO+CD69+CD103+ T_RM_ signature were found in the synovium of all RA donors.

### Memory T cells within RA synovium exhibit gene expression consistent with T_RM_

To test whether these RA synovial T cells are T_RM_, we performed single-cell RNA sequencing (scRNA-seq). Leukocytes were harvested from disaggregated synovial tissue from five human RA donors (Supplemental Figure 5A). Memory T cells (CD3+CD45RO+) were enriched by fluorescence-activated cell sorting (FACS). Cells were clustered into a tSNE plot based on gene expression profiles (Figure 7C). Cells from all five donors were represented in each cell cluster, suggesting that clustering was robust to donor-to-donor variability (Supplemental Figure 5B-C).

Expression of published T_RM_-associated genes was evaluated within each cluster (Figure 7D) (6–8, 30). Transcription factors such as RUNX3, Hobit (*ZNF683*), or Blimp (*PRDM1*) are essential to T_RM_ differentiation (31, 32). CD49a (*ITGA1*), CD103 (*ITGAE*) and E-cadherin (*CDH1*) are cell adhesion molecules upregulated to anchor T_RM_ in tissue (9, 33, 34). T_RM_ downregulate receptors S1PR1 and CCR7 that facilitate emigration of lymphoid cells from tissue while expressing tissue retention mediators regulator of G-protein signaling-1 (*RGS1*) and *RGS2* (35–38). Only Clusters 3 and 4 displayed upregulation of at least one T_RM_-associated transcription factor, downregulation of *S1PR1* and *CCR7*, upregulation of either *ITGA1, ITGAE* or *CDH1*, and upregulation of *RGS1* and *RGS2* (Figure 7D). We performed gene set enrichment analysis comparing the gene expression within these clusters to the established T_RM_ transcriptional signature identified by Kumar et al. (30). Only Cluster 3 overlapped significantly with the lung T_RM_ transcriptome (Figure 7E). Of 1263 cells in Cluster 3, 1043 expressed detectable mRNA for *CD8a* and 96 for *CD4*, identifying 83-92% of these cells as CD8+, a proportion similar to the 10:1 CD8:CD4 ratio in T_RM_ defined by immunofluorescence. These support the identification of Cluster 3 synovial memory T cells as bona fide T_RM_.

### RA synovial T_RM_ express genes involved in inflammation and cellular recruitment

Gene ontology analysis of differentially expressed (DE) genes within Cluster 3 compared to all other clusters showed enrichment of several functional categories, including immune cell activation, cytokine and chemokine signaling, and other immune responses (Supplemental Figure 6A). Furthermore, Ingenuity Pathway Analysis revealed an enrichment of cellular recruitment pathways among genes upregulated within Cluster 3 and identified *CCL5* to be a hub gene within the gene network (Supplemental Figure 6B). This finding suggests that T_RM_ serve a pro-inflammatory role within the human arthritic joint, particularly through recruitment of infiltrating immune cells, operating via pathways similar to those employed by murine synovial T_RM_.

### Synovial T_RM_ signature widely identified in inflammatory arthritis, lymphocytic pathotype

To establish the general prevalence of T_RM_ in inflammatory arthritis, we assessed whole-synovium microarray expression data from two published studies (n=30 and n=15 samples) that compared RA tissues with healthy synovium and osteoarthritis (OA) synovium (39, 40). Using our Cluster 3 T_RM_ gene signature, we found that this signature was present in each of the RA samples but not the healthy or OA synovial tissues (Figure 7F-G).

We further interrogated whole-synovium RNA-seq from 66 well-characterized early-RA samples with varying RA pathotypes (fibroid, lymphoid, and myeloid) (41). Using either our Cluster 3 gene signature or the Kumar et al. signature, we found evidence for T_RM_-pattern gene expression in many samples (Supplemental Figure 7A-B). Intriguingly, but perhaps not surprisingly, the signature was most evident in the lymphoid subset, though a few donors within the myeloid and fibroid subset also exhibited a T_RM_ signal. No difference in T_RM_ signal was noted as a function of time between first symptom and sampling (all less than 12 months), consistent with the observation that disease duration in these early-arthritis samples was also not associated with pathotype (Supplemental Figure 7C-D). Using available clinical data, we found that the presence of T_RM_ correlated with systemic inflammation, reflecting the association of inflammation with the lymphoid pathotype (Supplemental Figure 7E-F). These confirm the widespread presence of T_RM_ in RA, particularly in inflamed synovium with a lymphocytic pathotype.

### T_RM_ gene signature is enriched in late-stage non-inflamed RA synovium

To determine whether T_RM_ are present in RA synovium during remission, we assessed bulk RNA-seq data from T cells sorted from 34 RA synovial tissue biopsies across a spectrum of synovial inflammation. Zhang et al. classified these synovial tissues as leukocyte rich, with T and B lymphocyte infiltration, or leukocyte poor, showing low inflammation comparable to OA tissues (42). Using our Cluster 3 gene signature, we found T_RM_-pattern gene expression to be relatively enriched in synovial tissues that were non-inflamed (leukocyte poor) compared to those that were inflamed (leukocyte rich), consistent with retention of T_RM_ cells in the synovium after inflammation has resolved (Figure 7H). Comparing Cluster 3 gene signature in RA synovial tissues obtained early in disease (disease duration less than 12 months) to late disease without synovitis on histopathology (disease duration over 5 years with Krenn inflammation score ≤1) (Figure 7I), we found the T_RM_ signature to be enriched in late-stage noninflamed tissues, again supporting cell retention in chronic RA joints during remission.

### RA synovial T_RM_ exhibit a restricted T cell receptor repertoire

We had found murine T_RM_ to be activated by specific antigen. To assess comparable antigen-restriction in human T_RM_, we assessed T cell receptor (TCR) repertoire. The third complementarity determining region (CDR3) is the hypervariable region on each alpha and beta TCR chain responsible for recognizing processed antigen peptides. We sequenced the CDR3 region of paired alpha and beta TCR chains for each cell and plotted the results by frequency (Figure 7J). The vast majority of memory T cells showed unique TCR sequences, but 0.5% of clonotypes included at least 5 cells bearing the same CDR3 TCR, representing expanded clones. Notably, clonal T cells largely mapped to Cluster 3 (Figure 7K), the cluster exhibiting T_RM_ gene expression, across multiple donors (Supplemental Figure 5D). T cells within Cluster 3 showed the lowest TCR diversity normalized to the number of T cells (clonality index) (Figure 7L), confirming repertoire restriction and supporting a role for antigen-specific T_RM_ in arthritis recurrence, as observed in the murine system.

## Discussion

RA is an autoimmune diseases characterized by recurrent joint inflammation (2). Despite the systemic nature of this disease, arthritis flares commonly affect only a subset of joints, typically in a recurrent manner, implying the existence of a mechanism of local memory that has remained undefined (4). Here, we show that resident memory T cells (T_RM_) mediate joint-specific memory in arthritis, driving site-specific flares and targetable to ameliorate disease recurrence.

T cell studies in RA have focused primarily on CD4+ T cells because anti-citrullinated protein antibodies positive (ACPA+) RA is strongly associated with the MHC class II HLA-DRB1 alleles; however, RA also exhibits genetic associations with alleles in the MHC class I HLA-B locus, highlighting the likely importance of CD8+ T cells (43, 44). While CD8+ T cells are described in the synovium, their role in RA disease has not been further defined.

We found that CD8+ rather than CD4+ T_RM_ play a predominant role in arthritis recurrence. These cells were triggered via antigen, since neither non-specific activating signals nor circulating antigen-specific T cells could replicate their role in initiating disease flare. Together with our observation that human synovial T_RM_ employ a restricted TCR repertoire, with most of the 0.5% most abundant TCR clones residing within Cluster 3, these findings identify antigen exposure as the mechanism by which T_RM_ are activated in the context of joint flare, a finding consistent with a recent study using Nur77 reporter mice to establish T cell receptor-driven activation of T_RM_ in lung (45).

T_RM_ are defined by their capacity for long-term persistence in tissues, mediated through resistance to tissue egress and therefore to recirculation via the blood and lymphatics (6, 9). Parabiosis studies have been employed to demonstrate tissue residency in skin. When the vascular systems of two animals are joined, circulating leukocytes establish equilibrium between animals while tissue-resident cells, including T_RM_, do not (9). In our experimental system, we achieved a comparable demonstration by studying contralateral joints within a single animal. To show that T_RM_ do not traverse to other sites, we labelled synovial cells in one joint during remission and found that labelled T cells did not appear within other joints. Synovial T_RM_ also failed to migrate to tissue egress signals in vitro and exhibited preferential uptake of fatty acids consistent with the known metabolic skew of T_RM_ toward oxidative phosphorylation. T_RM_ persisted in joints for more than 8 months, triggering an arthritis flare in response to systemic antigen stimulation even after extended remission. Thus, the synovial T_RM_ we identify here are bona fide tissue resident memory T cells by virtue of surface signature, tissue retention, and metabolic features, providing the immunologic substrate for long-term autoimmunity in the joint (5, 6, 18).

In human RA synovium, we identified T_RM_ by cell surface markers as well as gene expression profile. By immunofluorescence microscopy, we find that CD8+CD69+CD103+ T_RM_ represent 1% of memory T cells in RA synovium, outnumbering CD4+CD69+CD103+ tenfold. However, these figures likely underestimate the T_RM_ population, as T_RM_ do not always express CD103 and display considerable heterogeneity between tissues (7, 8, 46, 47). Using scRNA-seq, cells with a T_RM_ gene expression signature represents up to 15% of synovial memory T cells (Cluster 3). Consistent with our immunofluorescence results, more than 80% of these cells are CD8+, establishing CD8+ T_RM_ as the most common resident memory population within RA synovium. The discrepancy in T_RM_ abundance between immunofluorescence and RNA-seq likely reflects the overly conservative nature of the T_RM_ surface signature we used for imaging (6). These findings in human RA synovium complement our murine observations in the importance of CD8+ rather than CD4+ T_RM_.

To understand how T_RM_ contribute functionally to arthritis flares, we show that secondary recruitment of circulating lymphocytes, particularly through the production of CCL5, is necessary for arthritis flare to occur in mice. In human RA, gene ontology analysis of differentially expressed genes showed that cells in Cluster 3 exhibit immune effector activity, supporting the hypothesis that T_RM_ play an active role in synovial inflammation within RA. Furthermore, Ingenuity Pathway Analysis of Cluster 3 differentially expressed genes shows an enrichment and upregulation of cellular recruitment pathways with CCL5 as a key hub gene within the network.

We developed a system to deplete resident T cells from within the joint during remission and found that local depletion impedes arthritis flare, demonstrating an essential role of T_RM_ within quiescent joints in instigating joint-specific flares. We highlight that a significant reduction in inflammation could be achieved with only partial T cell depletion (limited experimentally here by the maximum dose of intraarticular DT that did not also deplete circulating T cells), suggesting that more complete elimination of synovial T_RM_ via other methods could further attenuate or even abort recurrent disease. Thus, targeting synovial T_RM_ represents a novel therapeutic approach to disease recurrence in arthritis.

No animal model exactly mimics human disease. Thus, it is important to consider when murine data inform the understanding of human pathology. Perhaps most compelling is when orthogonal model systems reach the same conclusion. Here, we characterized T_RM_ in models of arthritis driven by antigen injection into the joint followed by systemic antigen challenge (meBSA and OVA + OT-I T cell transfer models), both in the C57BL/6 background. We find the same biology in a third model wherein Balb/c mice lacking endogenous IL-1Ra spontaneously develop chronic joint inflammation driven by T cells, IL-1, and IL-17 (27). Thus, T_RM_ are not limited to one animal model or to arthritis induced via an exogenous antigen, confirming the generalizability of T_RM_ in arthritis pathogenesis. Further, as in human disease, disease flares in our models were joint-restricted, and T_RM_ cells were predominantly CD8+ and antigen-specific, findings echoing those in human synovium.

Importantly, we could identify the T_RM_ gene signature in more than 100 RA synovial tissue samples. In transcriptome data from 45 synovial tissue biopsies published by Woetzel et al. and Ungethuem et al. (39, 40), we find that the T_RM_ gene signature is present in RA synovium but not osteoarthritis and healthy control tissue. In 66 early RA synovial tissue samples from diverse synovial pathotypes (41), the T_RM_ gene signature is preferentially found in RA synovium with lymphoid-type histology. In 34 RA synovial tissue biopsies across a spectrum of synovial inflammation (42), the T_RM_ gene signature is enriched in late-stage non-inflamed synovium compared to early and inflamed synovium, suggesting relative retention of T_RM_ in RA synovium during remission. These data establish the generalizability of our murine T_RM_ findings, human scRNA-seq data, and immunofluorescence imaging observations across human RA.

These findings represent the first confirmation that T_RM_ occur in arthritic synovium and play a direct role in disease pathogenesis. Cells with a related phenotype have been identified in arthritic synovial fluid. Petrelli et al. described a clonally expanded subset of PD-1+ expressing CD8+ T cells within inflamed synovial fluid expressing a higher level of the T_RM_ markers CD69 and CD103 than their PD-1-counterparts (48). Quayum et al. found a population of CD103+β7+CD29+CD49a+ “integrin-expressing” memory CD8+ T cells reminiscent of T_RM_ (CD62L-S1PR1-) in the synovial fluid of patients with ankylosing spondylitis, as well as in gut, suggesting a possible gut-joint connection (46, 49). It remains unclear whether the cells identified by these groups persist between flares to provide tissue-specific memory.

While our results establish T_RM_ as a mechanism of joint-specific memory, other mechanisms may also exist. Synovial fibroblasts represent a key effector lineage in arthritis, acquiring an invasive and pro-inflammatory character through epigenetic reprogramming (50–52). Synovial macrophages could potentially acquire a pathogenic character that persists even after a primary immune stimulus resolves (53). In arthritis mediated by autoantibodies, joint inflammation could recur through deposition of circulating antibodies or by autoantibodies generated locally by B cells with the assistance of tissue-resident T helper populations (2, 54). Depletion of synovial T_RM_ is thus likely to halt recurrent inflammation in some contexts but not others. Therapeutic implications across the range of human arthritides remain to be established (55).

Taken together, our data show that T_RM_ form in previously inflamed joints and persist in the synovium during remission. Activation of T_RM_ leads to arthritis flares with joint swelling and infiltration of circulating leukocytes, ultimately contributing to joint destruction (Supplemental Figure 8). While patients with RA currently require lifelong treatment (3), our findings imply that targeting local T_RM_ could provide a new option to achieve durable disease control without systemic immunosuppression.

## Methods

### Human Subjects

Synovial tissues were obtained from joint replacement surgeries, synovectomy surgeries or synovial biopsies of RA patients. The subjects were enrolled at Brigham and Women’s Hospital (BWH) with appropriate informed consent as required. The study protocols were approved by the Institutional Review Board of BWH.

### Human synovium processing

Formalin-fixed paraffin embedded (FFPE) synovial tissues used for microscopy were obtained from the pathology archives at BWH as unstained tissue sections mounted on slides. Fresh synovial tissues were obtained immediately after surgeries. Bone and adipose tissues were removed with scissors and the remaining synovium was cut into small pieces then subjected to enzymatic digestion and mechanical dissociation. Tissue fragments were homogenized with gentleMACS (Miltenyi) and digested in RPMI with Liberase TL 0.1mg/mL (Roche) and DNase I 0.1mg/mL (Sigma-Aldrich) for 30 minutes at 37°C before filtering through a 70μm cell strainer.

### Mantra microscopy

Opal Multiplex IHC Detection Kit was used to stain synovial tissues per the manufacturer’s directions (PerkinElmer). Briefly, unstained FFPE slides of synovial tissues were deparaffinized by heating in a dry oven at 58°C for 45 minutes, washed with xylene three times for a total of 40 minutes, then hydrated through an ethanol gradient ending with a distilled water wash. The slides were then fixed with 10% neutral buffered formalin for 10 minutes. Antigen retrieval was achieved by microwaving the slides in Tris-EDTA Buffer (10mM Tris, 1mM EDTA, pH 9.0) for 15 minutes. Slides were cooled then washed with TBS-Tween20 for 20 minutes, then blocked with BLOXALL (Vector Laboratories). Multiplex antibody staining was conducted with serial cycles of antigen retrieval, 1 hour incubation with primary antibodies against each target antigen, 10 minute incubation with Opal polymer HRP-labeled secondary antibodies, and 10 minute Opal fluorophore incubation for each fluorescent label. The slides were then mounted with media containing DAPI counter staining. Synovial sections were imaged with the Mantra Quantitative Pathology Workstation microscope platform by PerkinElmer. Images were captured using the Mantra Snap program version 1.0.4 at 20x magnification. 5-7 image fields containing CD3+ cells were chosen for each sample. Images were analyzed using the InForm software program version 2.4.8. Individual cells were segmented by identifying the nucleus of each cell through positive DAPI staining, cell membrane size was set to standard distance from the nucleus. Threshold for positivity for each fluorescent marker was generated automatically per image and confirmed manually. Positive expression of each cellular marker was defined by mean fluorescence intensity above the threshold for positivity for each individual cell. Coexpression was defined by positive expression of multiple cellular markers within the same cell.

### Single cell RNA sequencing

Viable cells were enriched by flow cytometric sorting using BD FACSAria Fusion and resuspended in 0.4% BSA in PBS at a concentration of 1,000 cells/μl. 12,000-15,000 cells were loaded onto a single lane (Chromium chip, 10X Genomics) followed by encapsulation in lipid droplets. cDNA and library generation per manufacturer protocol (Chromium Next GEM Single Cell 5’ Library & Gel Bead Kit v1.1, 10X Genomics). Additional target amplification for immune cell profiling were all prepared as per manufacturer protocol (Chromium Single Cell V(D)J Human T Cell Enrichment Kit, 10X Genomics). cDNA libraries were sequenced to an average of 50,000 reads per cell using Illumina Nextseq 500. scRNA-seq reads were processed with Cell Ranger v3.1, which demultiplexed cells from different samples and quantified transcript counts per putative cell. Quantification was performed using the STAR aligner against the GRCh38 version of the genome. scRNA-seq data was normalized using Cell Ranger. tSNE plots and clusters were calculated with Cell Ranger. Data were visualized using the Loupe Browser.

Differentially expressed genes between clusters were identified using Cell Ranger with thresholds FDR <0.1 and log2 fold change >0.5. We then compared these differentially expressed genes to genes identified as signatures of CD8 and CD4 T_RM_ from the lung as described in Kumar et al. (30). The significance of the overlap between each comparison was calculated using a hypergeometric probability.

Gene ontology was performed using DAVID (56, 57). Differentially expressed genes were input into DAVID, and GO_FAT categories Biological Process, Cellular Compartment, and Molecular Function were selected. Significantly enriched, non-redundant terms were selected. Ingenuity Pathway Analysis (IPA) (version 57662101) was used for pathway enrichment analysis in Ingenuity knowledge base (www.qiagen.com/ingenuity). Differentially expressed genes within Cluster 3 (p<0.01) were included in the analysis.

Gene set enrichment analysis was performed using the GSEA program. Briefly, differentially expressed genes that were upregulated within cluster 3 compared to all other clusters were compared to genes that were upregulated in CD8+ T_RM_s from the lung (30), which were pre-ranked based on fold change. The analysis was run without weighting based on ranking.

Gene expression data from additional synovial tissue specimens were obtained from publicly available datasets (ArrayExpress E-MTAB-6141(41); ImmPort SDY998(42); GEO DataSets GSE55235(39); GEO DataSets GSE1919(40)). Bulk RNA-seq data from different RA pathotypes (41) were aligned using STAR (2.7.0f) and quantified using the HTSeq-count package (0.11.1) in python using all protein coding Ensembl genes as a reference. Processed RNA-seq data was obtained from sorted RA synovial T cells (42) and microarray data from healthy, OA and RA synovial tissue (39) and (40). Differentially expressed genes from cluster 3 were identified in each dataset, and the principal components analysis was performed using the singular value decomposition algorithm on scaled, centered expression data.

For donor differentiation using SNPs, scRNA-seq data was examined for the top 100 highest expressed genes and reads mapping to these genes were extracted from mapped SAM files using samtools. SNPs present in the coding regions of these genes were extracted from the UCSC Genome Browser, and SNPs were limited to single nucleotide variants. We then examined each read to determine whether the SNP was present in the read. Genotyped SNPs were collected, and a determination of the genotype was attempted based on the information. If only the major allele was present in a given cell, then the SNP for that cell was given a value of *g*(*x*) = 1 − (2 * [*MaAF*] * [*MiAF*])^*N*^, where MaAF is the major allele frequency, MiAF is the minor allele frequency, and N is the number of reads containing the SNP. If both major and minor alleles were present, then the SNP was given a value of 0.5. If only minor alleles were present, then the SNP was given a value of *g*(*x*) = 0.5 − (*MiAF*)^*N*^. A matrix of all genotyped SNPs by all cells was created, and SNPs were limited by those with the greatest variance. Hierarchical clustering was performed, and the clusters were selected based on the number of donors.

### Immune cell profiling

The 10x Genomics Single Cell Immune Profiling platform was used for immune cell profiling. This technology integrates a step into the 10x single cell RNA sequencing workflow for targeted amplification of full-length VDJ sequences with primers for the T cell receptor CDR3 of both alpha and beta chains (Chromium Single Cell V(D)J Human T Cell Enrichment Kit, 10X Genomics). Cell Ranger was used to associated cell barcodes with specific TCR clonotypes. Shannon entropy was calculated 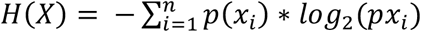 where p(xi) is the fraction of a single clonotype compared to all clonotypes present, and it was corrected for cluster size by *H*(*X*)/log_2_(*cells in cluster*).

### Murine arthritis models

#### Inducible arthritis model

Arthritis was engendered in 8-12 week old C57BL/6J (B6) mice (B6, JAX strain 664) or B6.SJL-Ptprc^a^Pepc^b^/BoyJ mice (B6 CD45.1, JAX strain 2014) by intraarticular injection of 400ug/10uL meBSA (Sigma-Aldrich) antigen into wrist, knee and ankle joints of a B6 mouse while the contralateral joints were injected with 10uL of PBS as an internal control. The mice also received 250ng/20μL of recombinant human IL-1β (Peprotech) into either footpad on Days 0-2 (20, 22). Arthritis flare was triggered by intraperitoneal injection of 400μg of meBSA. For adoptive transfer experiments, peripheral lymph nodes (cervical, brachial, inguinal, popliteal) were collected from 6 week old C57BL/6-Tg(TcraTcrb)110Mjb/J mice (OT-I, JAX strain 3831) or B6.Cg-Tg(TcraTcrb)425Cbn/J mice (OT-II, JAX strain 4194). Lymph nodes were disaggregated by mechanical dissociation over a 70μm cell strainer. T cells were purified by negative selection with the EasySep Mouse T cell isolation kit (STEMCELL Tech) as per manufacturer’s instructions. 2.5-5×10^6^ T cells were then transferred by retroorbital injection into B6 CD45.1 mice 24 hours before intraarticular injection of antigen. In adoptive transfer experiments, 500μg/10μL of OVA (Sigma-Aldrich) antigen was injected instead of meBSA. Intraperitoneal re-stimulation was conducted with 100μg of OVA.

#### Spontaneous arthritis model

*Il1rn-/-* mice were obtained from the Iwakura lab (27). These mice spontaneously develop arthritis with age. Remission was induced by daily injection of i.p. anakinra 20mg/kg (Sobi) for 15 days until there was no clinical signs of joint swelling. Arthritis flare was achieved by discontinuation of anakinra injections.

### Murine synovium processing

Synovium was dissected from the knees and ankles of mice and digested for 30 minutes with DNase I 0.1mg/mL (Sigma-Aldrich) and Collagenase type IV 1mg/mL (Worthington) in RPMI at 37°C. The tissue was then mechanically dissociated over a 70μm cell strainer into a cell suspension for subsequent assays.

### Joint histopathology

The knees were dissected en bloc by amputation at mid-femur and mid-tibia. The skin, fat and muscles were trimmed off with scissors. The knee was fixed in 4% paraformaldehyde for 24 hours, protected from light at 4°C, then washed with PBS and placed into Osteosoft (Sigma-Aldrich) decalcification solution for 3-5 days. The tissue was embedded in paraffin, sectioned and stained with hematoxylin and eosin for light microscopy. Inflammatory score calculated as previously described (58).

### In vivo cell tracking

Arthritis was engendered in 8-12 week old B6.129(Cg)-Gt(ROSA)26Sor^tm4(ACTB-tdTomato--EGFP)Luo^/J mice (mT/mG, JAX strain 7676) with intraarticular antigen injection as described above. During remission, 5μL of adeno-associated virus serotype 2/6 expressing Cre recombinase (3.88×10^12^ genome copies/mL) was intraarticularly injected into one knee while 5μL of PBS control was injected into the contralateral knee. 4 weeks later, knee synovium was collected and processed for analysis.

### Localized T cell depletion in vivo

B6.Cg-Tg(Lck-cre)548Jxm/J mice (Lck-Cre, JAX strain 3802) were crossed with C57BL/6-Gt(ROSA)26Sor^tm(HBGEF)Awai^/J mice (B6-iDTR, JAX strain 7900) to generate mice with *Lck-driven* diphtheria toxin receptor expression (Lck-iDTR). These mice express the diphtheria toxin receptor on all T cells, rendering them susceptible to diphtheria toxin (DT)-mediated depletion. Arthritis was engendered in both knees of 8-12 week old Lck-iDTR mice with intraarticular antigen injection as described above. During remission, 25ng of DT (Sigma-Aldrich) was injected into one knee while the contralateral knee received intraarticular PBS. Arthritis flare was triggered 2 weeks later with i.p. antigen challenge. Synovium was collected 72 hours later and assessed by flow cytometry as described above.

### T cell migration

T cells were isolated from disaggregated synovium with the EasySep Mouse T cell isolation kit (STEMCELL Tech) as per manufacturer’s instructions. T cells were placed in the top chamber of a transwell polycarbonate membrane insert with a 5.0μm pore (Corning) in serum-free RPMI 1640 media. RPMI 1640 medium containing 10% FBS and varying concentrations of CCL21 (Peprotech) was placed in the bottom of a 24-well plate. Cells were allowed to migrate for 2 hours at 37°C, after which cells were collected separately from the top and bottom chambers and stained for flow cytometry analysis.

### Fatty acid uptake

Cell isolated from synovium were incubated with 1μM Bodipy FL C16 (ThermoFisher) in PBS with 20μM fatty acid-free BSA (Sigma-Aldrich) for 30 minutes at 37°C. Bodipy uptake was quenched by adding 4x volume of ice-cold PBS with 2%FBS, then cells were washed twice before antibody staining for flow cytometry analysis. All antibody staining and washing was performed on ice.

### Antibodies

Anti-Thy1.2 antibody (Clone 30H12, Bio X Cell Cat# BE0066, RRID:AB) was used for the depletion of circulating lymphocytes. Rabbit polyclonal anti-CCL5 antibody (Origene Cat# PP1081P1, RRID:AB_1006884) was used for the neutralization of CCL5 chemokine.

The following antibodies were used for the analysis of human synovial cells with cell sorting: APC anti-CD3 (Clone OKT3, BioLegend Cat# 317317, RRID:AB_1937213), PE anti-CD45RO (Clone UCHL1, BioLegend Cat# 304206, RRID:AB_314422).

The following antibodies were used for the analysis of human synovium with microscopy: rabbit polyclonal anti-CD3 (Sigma-Aldrich Cat# C7930, RRID:AB_259074), anti-CD103 (Clone EP206, Sigma-Aldrich Cat# 437R), rabbit polyclonal anti-CD69 (Sigma-Aldrich Cat# HPA050525), anti-CD4 (Clone EPR6855, Abcam Cat# ab133616), anti-CD8 (Clone C8/144B, BioLegend Cat# 372902), anti-CD45RO (Clone UCHL1, BioLegend Cat# 304202, RRID:AB_314418).

The following antibodies were used for the analysis of murine synovium with flow cytometry or cell sorting: Alexa Fluor 700^®^ anti-CD3 (Clone 17A2, BioLegend Cat# 100216, RRID:AB_493697), APC anti-CD3 (Clone 17A2, BioLegend Cat# 100236, RRID:AB_2561456), FITC anti-CD45 (Clone 30-F11, BD Pharmingen Cat# 561088), FITC anti-CD45.1 (Clone A20, BioLegend Cat# 110706, RRID:AB_313495), PE anti-CD45.2 (Clone 104, BioLegend Cat# 109808, RRID:AB_313445), FITC anti-CD4 (Clone GK15, BioLegend Cat# 100406, RRID:AB_312691), APC/Cyanine7 anti-CD4 (Clone GK1.5, BioLegend Cat# 100414, RRID:AB_312699), Brilliant Violet™ 650 anti-CD8a (Clone 53-6.7, (BioLegend Cat# 100742, RRID:AB_2563056), PerCP/Cyanine5.5 anti-CD8a (Clone 53-6.7, BioLegend Cat# 100734, RRID:AB_2075238), Brilliant Violet™ 605 anti-CD44 (Clone IM7, BioLegend Cat# 103047, RRID:AB_2562451), APC anti-CD62L (Clone MEL-14, BioLegend Cat# 104412, RRID:AB_313099), Pacific Blue™ anti-CD69 (Clone H1.2F3, BioLegend Cat# 104524, RRID:AB_2074979), PE/Cyanine7 anti-CD103 (Clone 2E7, BioLegend Cat# 121426, RRID:AB_2563691), PE anti-CD103 (Clone 2E7, BioLegend Cat# 121406, RRID:AB_1133989), PE anti-ST2 (Clone DIH4, BioLegend Cat# 146608, RRID:AB_2728177), PE anti-CD19 (Clone eBio1D3, Thermo Fisher Scientific Cat# 12-0193-82, RRID:AB_657659), APC anti-CD11b (Clone M1/70, BioLegend Cat# 101212, RRID:AB_312795), Brilliant Violet™ 650 anti-Ly6G (Clone 1A8, BioLegend Cat# 127641, RRID:AB_2565881), Brilliant Violet™ 421 anti-NK1.1 (Clone PK136, BioLegend Cat# 108731, RRID:AB_10895916). 123count eBeads (Invitrogen) were used to calculate absolute cell counts.

### Statistics

Statistical analyses were performed with GraphPad Prism software. Comparisons between contralateral joints employed two-tailed paired student’s t-test or two-tailed Wilcoxon matched-pairs signed rank test where indicated. Comparisons between two conditions utilized two-tailed student’s t-test or two-tailed Mann-Whitney test. Comparisons between three conditions were performed with one-way analysis of variance test. The RNA sequencing data were loaded into R and custom functions were used to analyze the data. Significance was determined using the tests described above.

### Data availability

The data that support the findings of this study are available on request from the corresponding author (PAN). Portions of the raw sequencing data will not be publicly available due to institutional restrictions regarding the potential to contain information that may compromise research participant privacy.

### Study Approval

Human subjects were enrolled at BWH with appropriate informed consent as required. Human study protocols were approved by the Institutional Review Board of BWH. Animal study protocols were approved by the Institutional Animal Care and Use Committee at BWH.

## Author Contributions

MHC, AL, RCF, PAN conceived and designed the study. KW, LAH, RAC, DAR provided expertise and assisted with the design of human studies. NNM, RBB, KDW, AM, SP, AW, RGB performed the experiments and acquired data. YI provided essential reagents. MHC, KDW analyzed the data. MHC, PAN drafted the manuscript and all authors edited the manuscript.

## Acknowledgements

MHC was supported by a Rheumatology Research Foundation Scientist Development Award, Boston Children’s Hospital Lovejoy Award, NIH/NIAID T32AI007512, NIH/NICHD K12HD052896, a Joint Biology Consortium microgrant off parent grant NIH/NIAMS P30AR070253, and a Translation Accelerator Grant from the Human Skin Disease Research NIH/NIAMS P30AR069625. AL was supported by a Joint Biology Consortium microgrant off parent grant NIH/NIAMS P30AR070253. KDW was supported by American Academy of Neurology Neuroscience Research Training Scholarship. KW was supported by NIH/NCATS KL2TR002542 and NIH/NIAMS P30AR070253. LAH was supported by NIH/NIAMS K08AR073339 and P30AR070253 an Investigator Award from the Rheumatology Research Foundation, and the Arthritis National Research Foundation. RAC was supported by NIH/NIAMS P30AR069625, NIH/NIAID R01AI127654, NIH/NIAIMS R01AR074797 and NIH/NCI R01CA203721. RCF was supported by an Innovative Research Grant from the Rheumatology Research Foundation and by NIH/NIAMS R01AR075906. PAN was funded by an Innovative Research Grant from the Rheumatology Research Foundation; NIH/NIAMS 2R01AR065538, R01AR075906, R01AR073201, P30AR070253, R21AR076630 and NIH/NHLBI R21HL150575; the Fundación Bechara; and the Arbuckle Family Fund for Arthritis Research.

