## Supplemental figures for "Arthritis Flares Mediated by Tissue Resident Memory T Cells in the Joint"

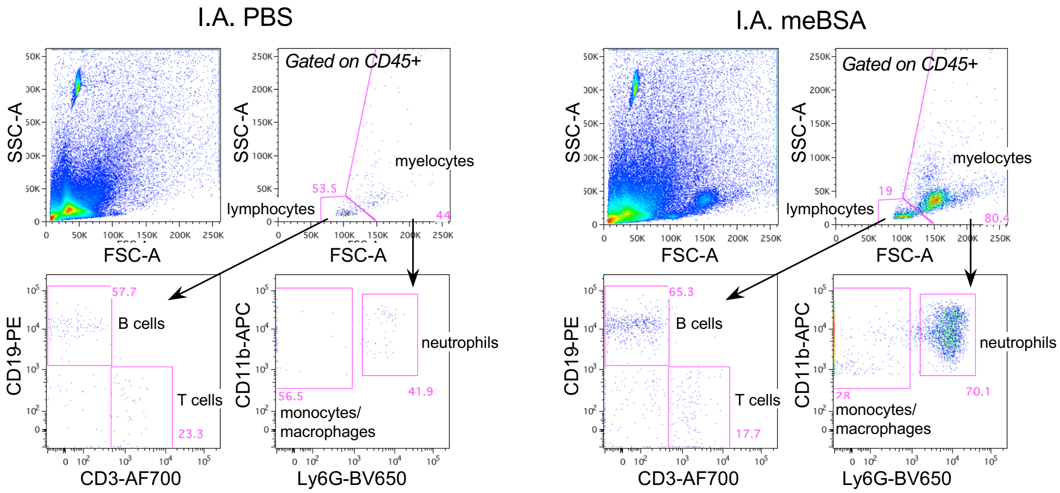


**Supplemental Figure 1. Lymphoid and myeloid cells are involved in arthritis flare.** Representative flow cytometry dot plot analysis of synovium on Day 31 flare. Left column depicts the joint injected with PBS control while the right column depicts the joint injected with meBSA antigen from the same mouse. The top left panel in each column shows the FSC vs. SSC of the whole disaggregated synovium. Top right panel shows gating of lymphocyte and myelocyte subpopulations based on FSC vs. SSC. Bottom left panel analyzes CD19 and CD3 expression in lymphocytes. Bottom right panel analyzes CD11b and Ly6G expression in myelocytes.

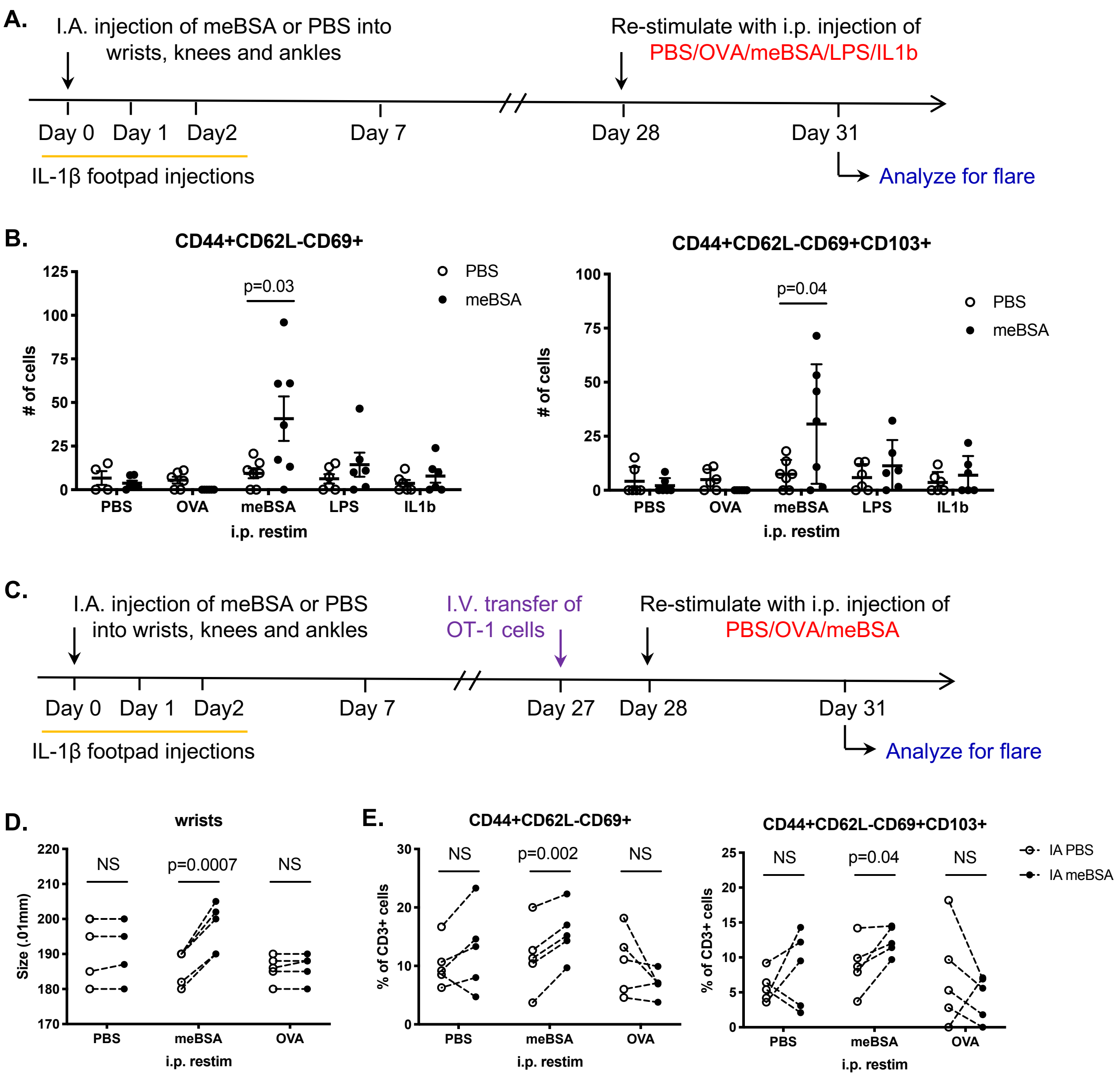


**Supplemental Figure 2. Arthritis flare in murine model is antigen specific. (A)** Experimental design for re-stimulating arthritis flare with nonspecific triggers. **(B)** Graphs depicting quantity of $T_{RM}$ in the synovium 72 hours after re-stimulation with PBS, OVA 250ug, meBSA 400ug, LPS 20ug or IL-1b 250ng. Each dot represents values for combined knee and ankle PBS- (open circle) and meBSA-injected (closed circle) joints from each animal (n=6 PBS, OVA, LPS, IL-1b re-stimulated mice; n=7 meBSA re-stimulated mice). p-values from two-tailed paired student's t-test. **(C)** Experimental design for re-stimulating arthritis with nonspecific antigen challenge. ***(D-E)*** Graphs depicting **(D)** wrist thickness and **(E)** prevalence of $T_{RM}$ in PBS- (open circle) vs. meBSA-injected (closed circle) joints at Day 31, flare. Dashed line connects contralateral joints within the same animal (n=5 mice per re-stimulation condition). p-values from two-tailed paired student's t-test.

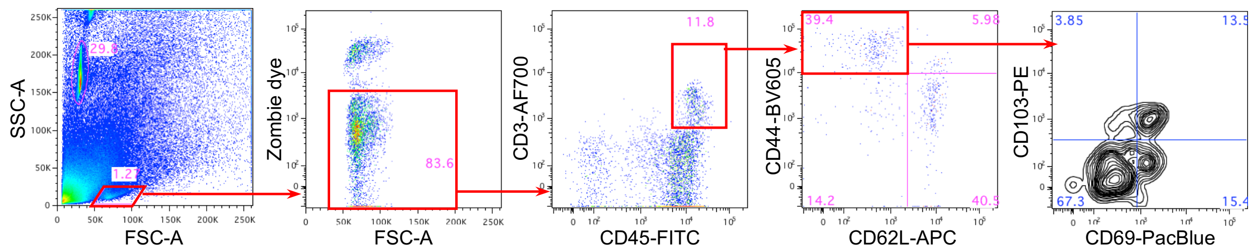


**Supplemental Figure 3. Gating strategy for $T_{RM}$ cells.** Representative flow cytometry plots of disaggregated synovial tissues depicting gating strategy used in data analysis. Left panel identifies lymphocyte population based on FSC vs. SSC. Second panel gates on Zombie dye negative live cells, followed by the identification of CD45+CD3+ T cells in the middle panel. Effector and resident memory T cells are then determined by CD44+ and CD62L- expression in the fourth panel, followed by analysis of CD69 and CD103 expression in the rightmost panel.

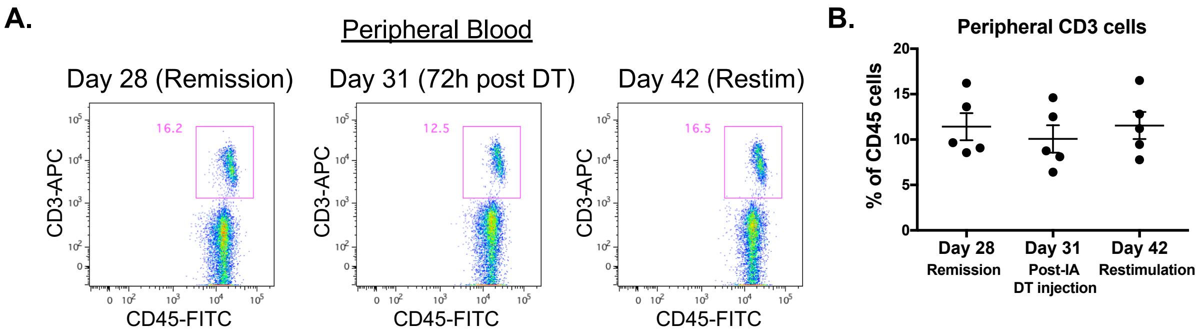


**Supplemental Figure 4. Local DT injection does not deplete circulating T cells. (A)** Representative dot plots taken from the same mouse showing percentage of T cells within peripheral blood before intraarticular DT injection (Day 28), 72 hours after a single DT injection, and at the time of re-stimulation, 2 weeks after DT injection (Day 42). **(B)** Graph showing percentage of peripheral blood T cells at different time points. Each dot represents one animal (n=5 mice).

**A.**

| Subject Characteristics | n=5 |
|---|---|
| Sex, female, n (%) | 4 (80%) |
| Age at time of biopsy, years, mean (range) | 66 (56-80) |
| Duration of disease, years, mean (range) | 12.4 (6-38) |
| RF+ or CCP+, n (%) | 2 (40%) |
| Surgery | |
| Arthoplasty | 4 (80%) |
| Arthrodesis | 1 (20%) |
| Medications at time of surgery | |
| non-biologic DMARD | 3 (60%) |
| biologic DMARD | 0 (0%) |
| prednisone | 3 (60%) |

**B.**

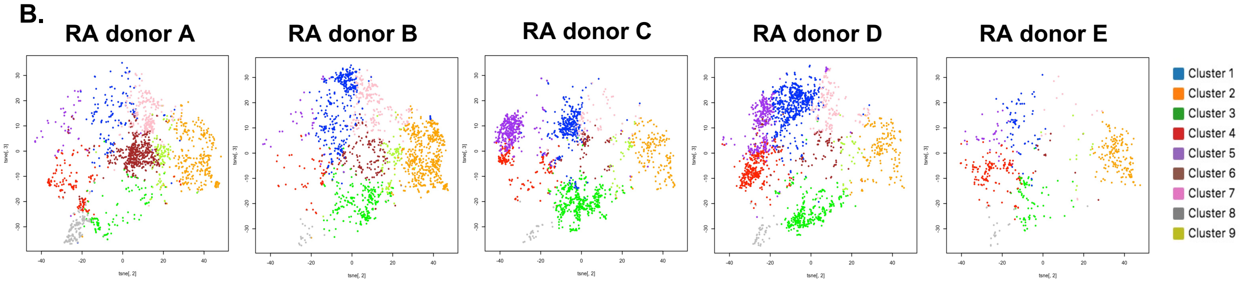


**C.**

| | Donors | | | | | |
|---|---|---|---|---|---|---|
| **Clusters:** | **RA A** | **RA B** | **RA C** | **RA D** | **RA E** | **Total cells** |
| **1** | 139 | 397 | 342 | 637 | 67 | 1582 |
| **2** | 314 | 581 | 176 | 168 | 171 | 1410 |
| **3** | 127 | 249 | 468 | 329 | 88 | 1261 |
| **4** | 188 | 55 | 184 | 352 | 157 | 936 |
| **5** | 50 | 38 | 421 | 349 | 36 | 894 |
| **6** | 510 | 146 | 73 | 103 | 15 | 847 |
| **7** | 261 | 215 | 75 | 190 | 15 | 756 |
| **8** | 189 | 20 | 32 | 45 | 25 | 311 |
| **9** | 118 | 95 | 42 | 35 | 13 | 303 |
| **Total cells** | 1896 | 1796 | 1813 | 2208 | 587 | 8300 |

**D.**

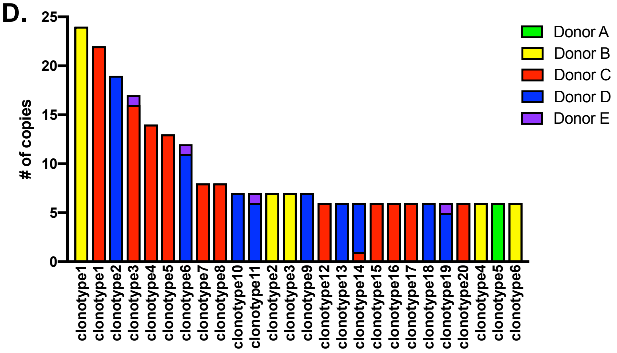


**Supplemental Figure 5. Characteristics and distribution of RA donors in single cell expression analysis. (A)** Clinical characteristics of human subjects. **(B)** Contribution of each donor to the tSNE plot of CD3+CD45RO+ cells from disaggregated RA synovium. Each dot represents one cell. **(C)** Table depicting the distribution of cells from each donor into the numbered cell clusters with similar gene expression profiles. **(D)** Top 0.5% of the most frequent T cell clones by donor. Each of the 5 individual RA synovium donors is designated with a different color (red, yellow, green, blue, violet).

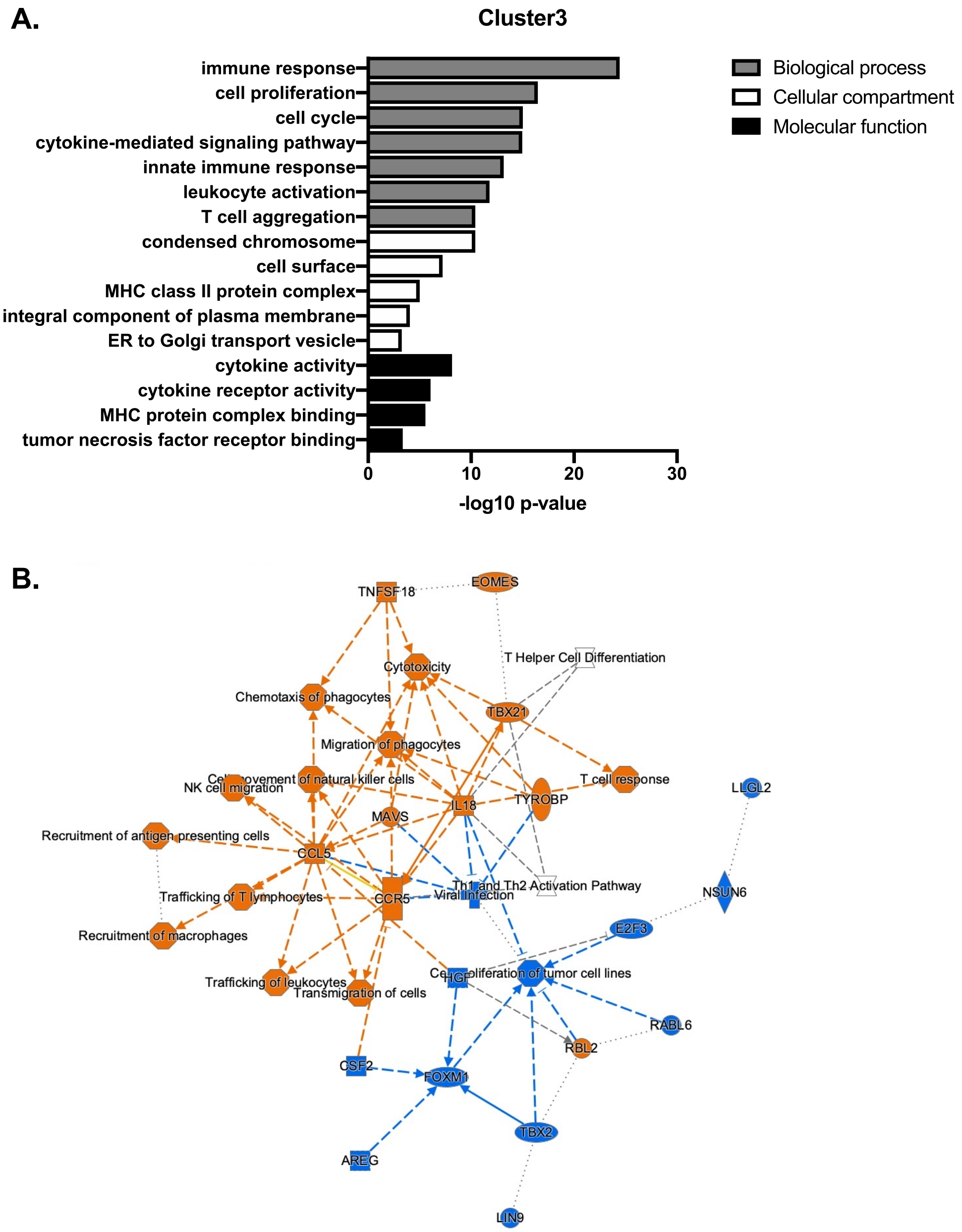


**Supplemental Figure 6. Functional analysis of Cluster 3 differentially expressed (DE) genes. (A)** Bar plot showing the log10 p-values of functional annotation of DE genes in Cluster 3 using gene ontology. **(B)** Graphical summary of Ingenuity Pathway Analysis of DE genes in Cluster 3. Orange represents pathways enriched among upregulated genes. Blue represents pathways enriched among downregulated genes.

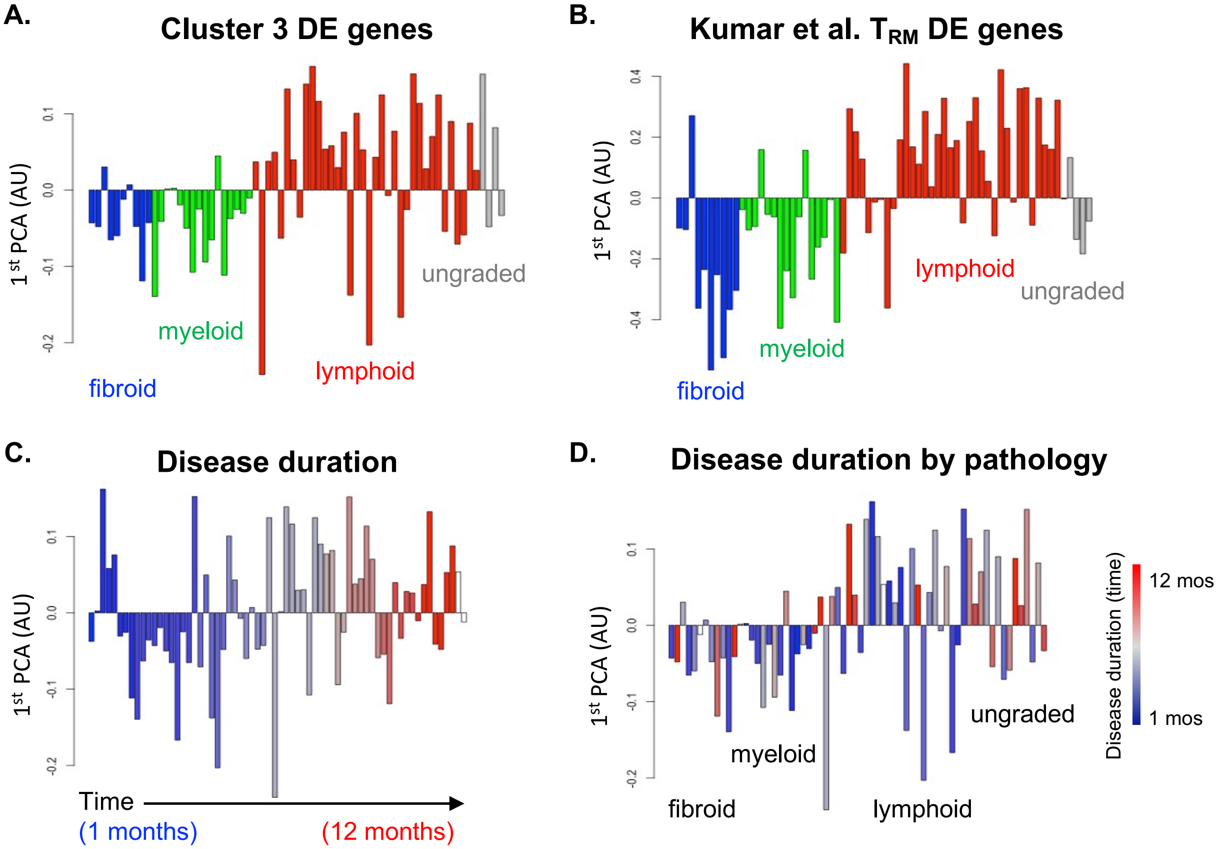


**E.**

| Factors | R-value | p-value |
|---|---|---|
| Age | -0.175 | 0.160 |
| Onset | 0.018 | 0.887 |
| ESR | 0.352 | 0.004 |
| CRP | 0.247 | 0.048 |
| CCP | 0.180 | 0.147 |
| RF | 0.154 | 0.217 |
| VAS | 0.236 | 0.056 |
| Tender | 0.190 | 0.127 |
| Swollen | 0.124 | 0.320 |
| HAQ | 0.254 | 0.041 |
| DAS28 | 0.309 | 0.012 |
| Inflammatory score | 0.641 | 4.54E-08 |

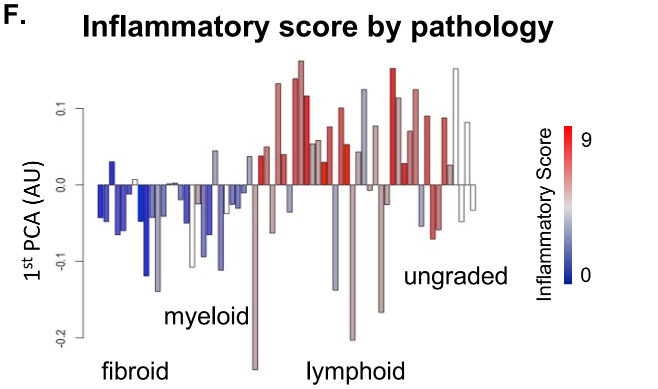


**Supplemental Figure 7. Cluster 3 $T_{RM}$ gene signature in early RA synovial bulk RNAseq data.** ***(A-B)*** Principal component analysis (PCA) of $T_{RM}$ differentially expressed (DE) genes in early RA synovial tissue. Bar plot shows the $T_{RM}$ gene expression signature identified from **(A)** Cluster 3 and **(B)** Kumar et al., 2017 in bulk RNAseq data from different RA pathotypes (fibroid, myeloid, lymphoid; n=66) published by Lewis et al., 2019. Each bar represents one donor. ***(C-D)*** Graph of 1$^{st}$ principal component of Cluster 3 $T_{RM}$ gene signature in RA synovial samples compared to **(C)** disease duration at time of biopsy or **(D)** disease duration within RA pathotypes. Each bar represents one donor. **(E)** Table outlining the correlation of clinical factors with the 1st principal component of Cluster 3 $T_{RM}$ gene signature. **(F)** Bar plot shows the histopathologic inflammatory score within RA pathotypes.

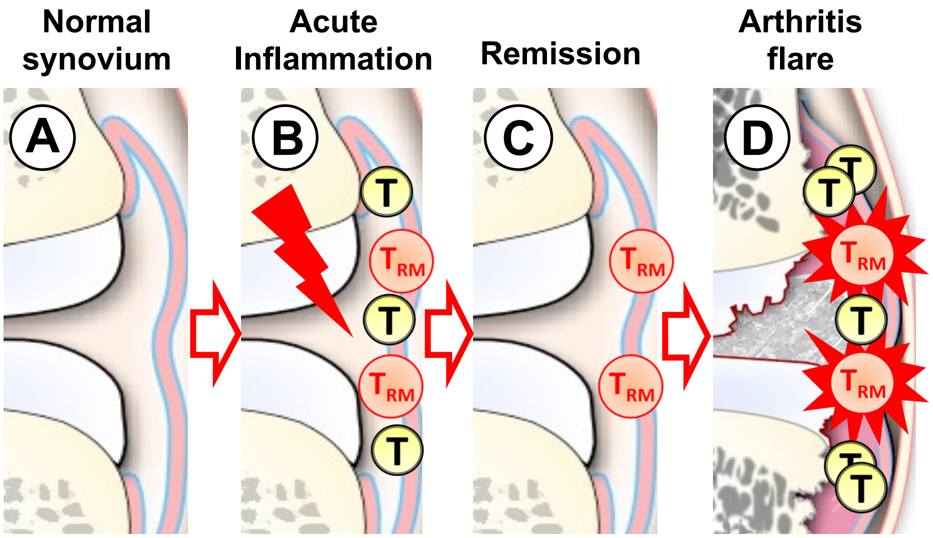


**Supplemental Figure 8. Model of synovial resident T cells mediating arthritis flare.** Illustration of how synovial TRM cells mediate arthritis flare. **(A)** Normal Synovium. **(B)** T cells are recruited to the joint during acute inflammation, some of which differentiate into TRM cells. **(C)** TRM cells remain in the synovium during remission after joint inflammation has resolved. **(D)** Activation of TRM cells triggers recurrent inflammation in the joint, leading to joint destruction.
